# Potent CRISPR-Cas9 inhibitors from *Staphylococcus* genomes

**DOI:** 10.1101/799403

**Authors:** Kyle E. Watters, Haridha Shivram, Christof Fellmann, Rachel J. Lew, Blake McMahon, Jennifer A. Doudna

## Abstract

Anti-CRISPRs (Acrs) are small proteins that inhibit the RNA-guided DNA targeting activity of CRISPR-Cas enzymes. Encoded by bacteriophage and phage-derived bacterial genes, Acrs prevent CRISPR-mediated inhibition of phage infection and can also block CRISPR-Cas-mediated genome editing in eukaryotic cells. To identify Acrs capable of inhibiting *Staphylococcus aureus* Cas9 (SauCas9), an alternative to the most commonly used genome editing protein *Streptococcus pyogenes* Cas9 (SpyCas9), we used both self-targeting CRISPR screening and guilt-by-association genomic search strategies. Here we describe three new potent inhibitors of SauCas9 that we name AcrIIA13, AcrIIA14 and AcrIIA15. These inhibitors share a conserved N-terminal sequence that is dispensable for anti-CRISPR function, and have divergent C-termini that are required in each case for selective inhibition of SauCas9-catalyzed DNA cleavage. In human cells, we observe robust and specific inhibition of SauCas9-induced genome editing by AcrIIA13 and moderate inhibition by AcrIIA14 and AcrIIA15. We also find that the conserved N-terminal domain of AcrIIA13-15 binds to an inverted repeat sequence in the promoter of these Acr genes, consistent with its predicted helix-turn-helix DNA binding structure. These data demonstrate an effective strategy for Acr discovery and establish AcrIIA13-15 as unique bifunctional inhibitors of SauCas9.

## Introduction

CRISPR systems are RNA-guided, adaptive immune systems that defend prokaryotes against invading mobile genetic elements (MGEs) (1). However, some MGEs, particularly phages, have evolved anti-CRISPRs (Acrs), peptide inhibitors of Cas proteins that block CRISPR defense systems (2, 3). Acrs have been discovered to inhibit distinct CRISPR systems, including type I (4-8), type II (9-16), type III (17, 18), and type V (7, 19). Strategies for identifying new Acrs include testing genes of unknown function that are proximal to anti-CRISPR associated (*aca*) genes (5-7, 9, 15) and screening genes in organisms with self-targeting CRISPR systems (7, 19, 20).

The frequent clustering of Acr and *aca* genes together allows for a ‘guilt-by-association’ approach that quickly identifies potential Acr candidates for experimental testing, but requires a known Acr or *aca* gene to seed the search (5-7, 9, 15). Conversely, self-targeting CRISPR systems represent diverse genomes that could encode corresponding CRISPR-Cas inhibitors to block autoimmunity (21) (Fig. 1A), but finding relevant Acr-encoding genes within these genomes can be daunting. Nonetheless, analysis of known Acrs suggests that self-targeting CRISPR-containing genomes are attractive to study because they often encode at least one corresponding Acr (19).

**Figure 1.**
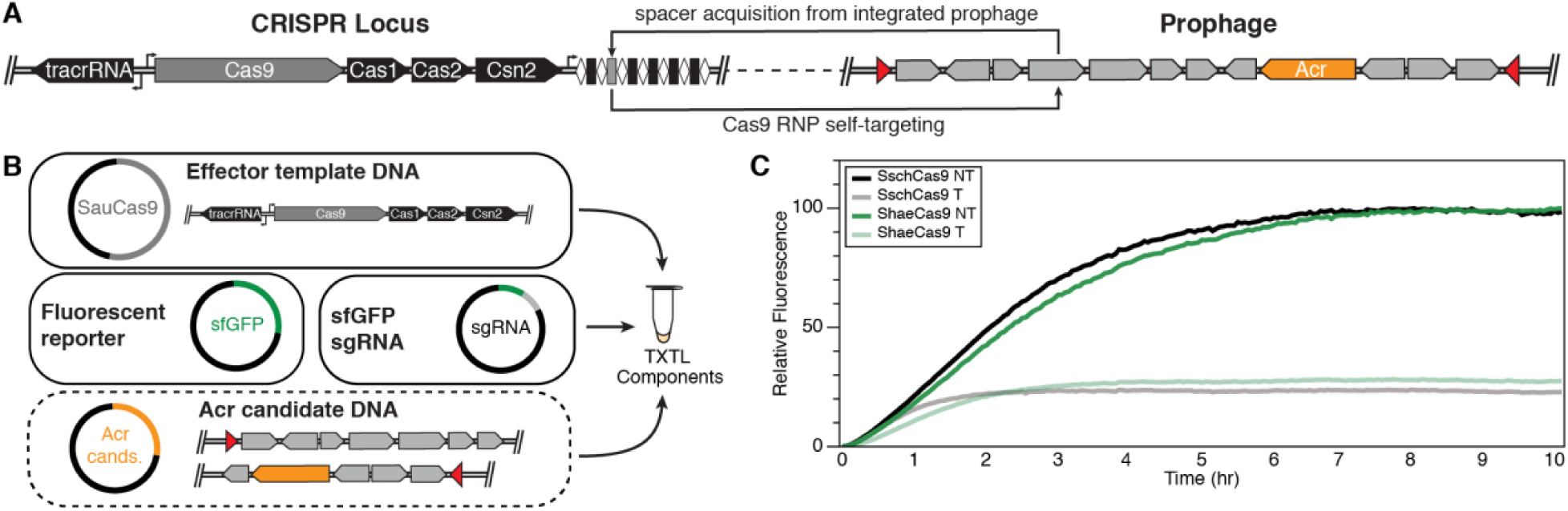
Identification of self-targeting Staphylococcus strains that contain active type II CRISPR-Cas systems. (A) Acquisition of protospacers from MGEs can result in self-targeting if the MGE is capable of stably integrating (e.g. prophages) or associating (e.g. plasmid) with the genome. (B) Overview of TXTL assay to test for Acr activity. Cas9 template DNA is combined with an sfGFP reporter plasmid and a plasmid expressing a single guide RNA (sgRNA) targeting the reporter. In the absence of Acr activity, expression of the Cas9 RNP suppresses sfGFP expression. In the presence of Acrs, the Cas9 RNP is inhibited, resulted in increased fluorescence. (C) Genomic amplicons from *S. schleiferi* and *S. haemolyticus* containing Cas9 lower sfGFP expression with an sfGFP-targeting sgRNA, demonstrating that the natural CRISPR loci are active.

Multiple anti-CRISPR (Acr) families inhibit *Streptococcus pyogenes* Cas9 (SpyCas9) and various Cas12a proteins, and can be used in cell-based experiments to control genome editing outcomes (7, 10, 11, 13, 14, 22, 23). Although weak cross-reactivity with other non-cognate Cas9 orthologs has been detected for a subset of these, we wondered whether more potent and selective inhibitors of particular Cas9 variants might exist in nature. To address this question, we focused on *Staphylococcus* genomes that might encode inhibitors of *Staphylococcus aureus* Cas9 (SauCas9), a genome editing alternative to SpyCas9 whose smaller size could offer advantages for delivery into mammalian cells (24, 25). We used a combination of self-targeting CRISPR screening and guilt-by-association genomic searches to discover three new peptide inhibitors of SauCas9. We show that these SauCas9 Acrs, AcrIIA13, AcrIIA14 and AcrIIA15, limit or prevent RNA-guided DNA cleavage *in vitro* and genome editing in human cells. These three inhibitors share a common N-terminal domain with a predicted helix-turn-helix structure that is dispensable for Acr function but can bind specifically to the inverted repeat sequence in the promoter of these Acr genes. The C-terminus of each Acr is distinct and is responsible for SauCas9 inhibition in each case, likely by differing mechanisms. These SauCas9 inhibitors provide new tools for the selective control of genome editing outcomes, and validate a multi-pronged strategy for discovering diverse Acrs in nature.

## Results

### Bioinformatic identification of self-targeting type II-A CRISPR systems

To identify potential new Acrs that inhibit SauCas9, we first used the Self-Target Spacer Searcher (STSS) (19) to query all *Staphylococcus* species deposited in NCBI for instances of CRISPR self-targeting. We observed 99 total instances of self-targeting in *Staphylococcus* CRISPR systems across 43 different strains out of a potential 11,910 assemblies searched (Table S1). Of the 99 self-targeting instances predicted, 50 could not be attributed to any particular CRISPR subtype, 48 were associated with a type II-A system, and one occurred as part of a type III-A system. We did not observe any self-targeting CRISPR type I-C systems that are occasionally found in *Staphylococcus* (26). It should also be noted that 29 of the predicted CRISPR self-targeting systems occurred in eight species whose CRISPR loci were manually annotated as type II-A based on identity to other type II-A Cas9-encoding genes.

To select the genomes most likely to contain Acrs, we filtered the list of 48 self-targets to exclude those with target protospacer-adjacent motifs (PAMs) that were more than one indel/mutation away from the known 3′-NNGRR(T) PAM for SauCas9 (25). This step eliminated genomes in which an incorrect PAM sequence could explain survival, without the need for Acrs. The remaining 14 self-targeting instances, belonging to 12 different strains (Table S1), were ranked according to similarity of their encoded Cas9 and SauCas9 (Table S2). Of the seven Cas9s displaying high similarity to SauCas9 (>50% identity with blastp), two species, *Staphylococcus haemolyticus* and *Staphylococcus schleiferi*, were selected for further screening for Acrs active against SauCas9 based on strain availability.

### Self-targeting screen for Acr activity

To reduce the sequence space to search for Acr genes, we selected only one of the two self-targeting strains identified for each species (*S. schleiferi* and *S. haemolyticus*), since the two strains of each respective species contained highly similar genomic sequences and MGE content. The *S. schleiferi* strain 5909-02 was chosen for having three self-targets vs. the two self-targets present in strain 2713-03, and *S. haemolyticus* strain W_75, containing a single self-target, was selected arbitrarily over strain W_139 (Fig. S1).

To first assess whether the CRISPR systems in *S. schleiferi* and *S. haemolyticus* are active, we generated amplicons encoding the Cas9, Cas1, Cas2 and Csn2 of each system under their natural promoters and a plasmid expressing a single-guide RNA (sgRNA) for SauCas9 (27). Two versions of the sgRNA were designed, one targeting a superfolder GFP reporter (sfGFP) and the other containing a non-targeting sequence. We co-expressed each amplicon with the targeting or control sgRNA-encoding plasmid, and a plasmid encoding sfGFP, in a cell-free transcription-translation extract (TXTL) (19, 28). We observed a strong reduction in sfGFP expression in the TXTL reaction mixtures expressing the targeting, but not the non-targeting, sgRNAs (Fig. 1C). This result suggests that the native type II-A CRISPR systems in both *S. schleiferi* strain 5909-02 and *S. haemolyticus* strain W_75 are active.

Nearly all currently known Acrs can be found in MGE sequences, especially phages, highlighting their use as a CRISPR counter-defense mechanism (3). Therefore, we examined the genomes of both *S. schleiferi* and *S. haemolyticus* for integrated MGEs using PHASTER (29) and Islander (30) to identify potential hotspots that might harbor Acrs. We found four predicted MGEs in *S. schleiferi* strain 5909-02 and eight in the contigs of the *S. haemolyticus* strain W_75 assembly. We then designed 27 genome fragment (GF) amplicons in total covering all the predicted sequences, each up to ∼10 kb, with the goal of best capturing complete operons wherever possible (Table S3). We then prepared TXTL reactions containing DNA encoding the sfGFP reporter, the sgRNA-encoding plasmid, a GF amplicon, and the associated Cas protein-encoding amplicon from the GF’s source organism. By comparing GFP expression over time, we found that the presence of GF5 from *S. schleiferi* corresponded to nearly complete inhibition of Cas9-mediated sfGFP knockdown (Fig. 2A). We observed no inhibitory activity in the presence of any GFs derived from *S. haemolyticus*.

**Figure 2.**
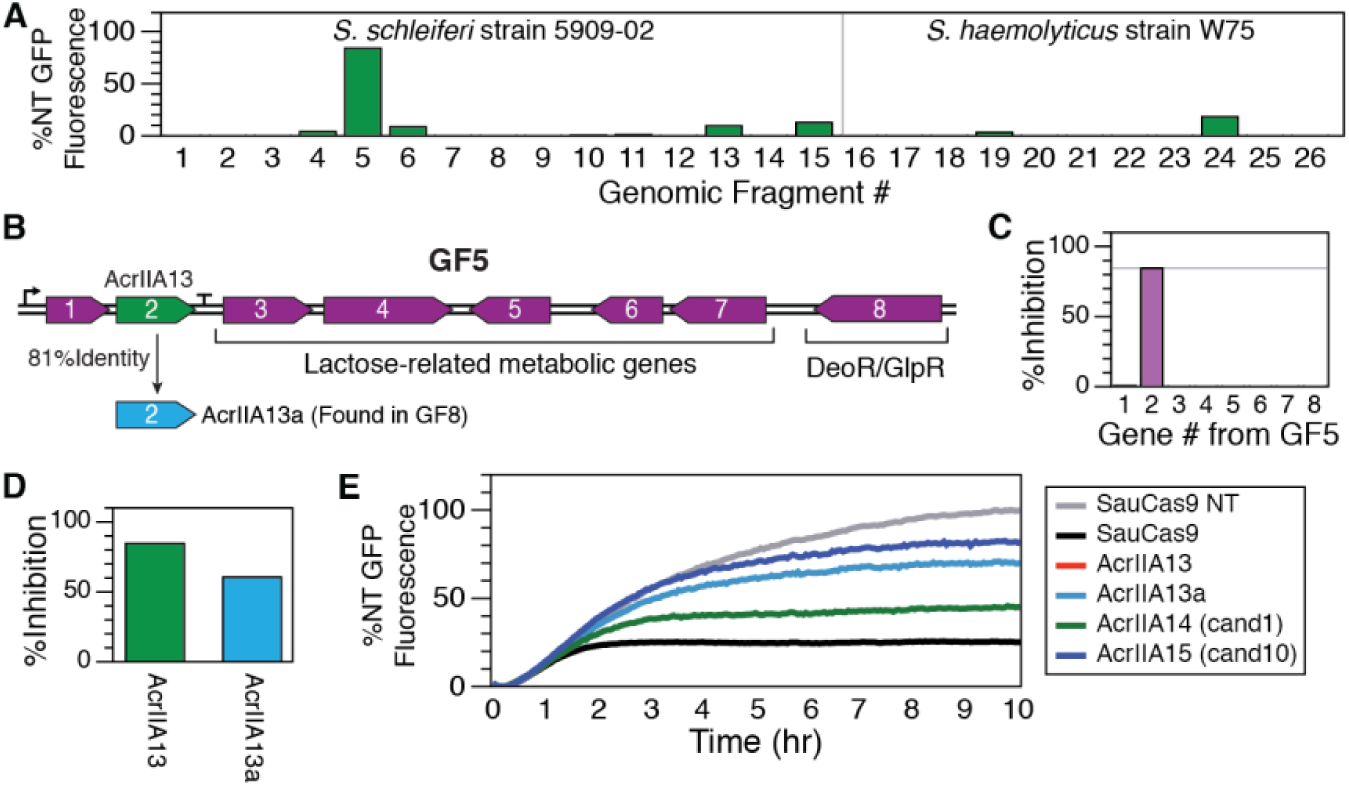
Identification of three new Acrs using amplicon screening and guilt-by-association. (A) The relative level of Cas9 DNA-cleavage inhibition for each fragment was measured as percentage of the GFP expression for the non-targeting (NT) control after subtracting the fluorescence level observed with no Acr present. Thus, 0% inhibition is equivalent to the GFP expression level measured with the targeting sgRNA, while 100% inhibition represents no reduction in the maximum GFP expression. GF5 is the only amplicon that exhibits Acr activity. (B) Genes found in GF5. (C) Each gene in GF5 was individually cloned and tested for Acr activity with TXTL and SauCas9. The 2^nd^ gene of GF5 (AcrIIA13) inhibits SauCas9. (D) AcrIIA13a, a homolog of AcrIIA13, is found in GF8 and is also able to inhibit SauCas9 in a TXTL assay. GF8, which contains AcrIIA13a did not exhibit Acr activity, however. (E) Of 10 candidates chosen from a guilt-by-association search seeded by GF5 gene 1 (*aca8*), candidates 1 (AcrIIA14) and 10 (AcrIIA15) were found to inhibit SauCas9 using the TXTL assay.

*S. schleiferi* GF5 contains eight genes (Fig. 2B), most of which are predicted to be part of the lactose metabolic pathway. Genes 1 and 2, however, have no predicted function. We cloned each gene from GF5 into a heterologous plasmid under the control of a pTet promoter and repeated the TXTL assay with each cloned gene individually in place of the GF5 amplicon. We observed that gene 2 alone produced the inhibitory effect previously observed with the complete GF5 (Fig. 2C). This gene was named AcrIIA13 to indicate its inhibition of a CRISPR type II-A system, in accord with standard Acr nomenclature (22). There is also a homologous protein, AcrIIA13a, found within GF8, and although no inhibitory activity from GF8 was ever observed, when AcrIIA13a is expressed separately using the pTet expression plasmid, it does exhibit inhibitory activity (Fig. 2D).

### ‘Guilt-by-association’ identification of SauCas9 Acrs

We wondered whether AcrIIA13 or the neighboring gene in its operon (GF5 gene 1) might be associated with other Acrs, and whether these inhibitors might have the ability to block activity of the SauCas9 protein. We first searched AcrIIA13 homologs to find potential Acr candidates co-occurring with it (Fig. S2A). While we did find a truncated homolog missing the AcrIIA13 N-terminus, AcrIIA13b, we did not observe any new genes commonly found near AcrIIA13. However, when we searched for homologs of GF5 gene 1, we found that they occur widely in bacteria (Fig. S2B), a typical hallmark of *aca* genes (6, 9, 10, 15). We identified genes that frequently co-occur with GF5 gene 1 and selected them as an initial set of Acr candidates. Then, because many Acrs are clustered together, we also added genes that co-occurred with the initially chosen Acr candidates to create the final test set. This set of ten candidate Acr genes was cloned into the pTet expression plasmid for further analysis.

Each of the ten new candidate Acr genes was tested in the TXTL sfGFP depletion experiment, this time using SauCas9 expressed from a recombinant plasmid. After ten hours, we observed that TXTL reaction mixtures for two of the new Acr candidates, candidate 1 and candidate 10, expressed GFP levels that were significantly above the baseline (Fig. 2E and S3). Candidate 1, hereafter referred to as AcrIIA14, is phylogenetically associated with a close homolog of GF5 gene 1 and occurs in *Staphylococcus simulans* strain 19, which had previously passed the PAM filtering step for self-targeting *Staphylococcus* genomes and has a Cas9 closely related to SauCas9 (86% identical via blastp) (Table S2). Although candidate 10, hereafter referred to as AcrIIA15, does not occur in a CRISPR-self-targeting strain, a close homolog (80% identical via blastp) can be found in *S. pseudintermedius* strain 104N, which is self-targeting (Fig. S2, Table S1).

### Reconstitution of Acr activity against SauCas9

To examine the inhibitory activity detected in the TXTL experiments with AcrIIA13-15, we purified each Acr protein and tested its ability to inhibit SauCas9-mediated double-stranded DNA (dsDNA) cleavage *in vitro*. We began by first assembling the SauCas9-sgRNA RNP complex, then added the individually purified Acr proteins and dsDNA target. Under these conditions, we observed near total inhibition of SauCas9 dsDNA cleavage at 20-fold molar excess of Acr relative to Cas9 RNP (Fig. 3A, left and Fig. S4A, top). As a negative control, we performed the same assay using BSA and AcrVA4 and observed no difference in cleavage activity of SauCas9 (Fig. S4B). To shed more light on the mechanism, we performed the SauCas9 *in vitro* cleavage experiments again, but added the Acr proteins before the addition of sgRNA. Adding the Acr protein before the sgRNA led to ∼4-fold increase in inhibition by the Acr proteins, on par with the only currently known broad spectrum Acr for type II-A Cas9, AcrIIA5 (11) (Fig. 3A right and Fig. S4A, bottom).

**Figure 3.**
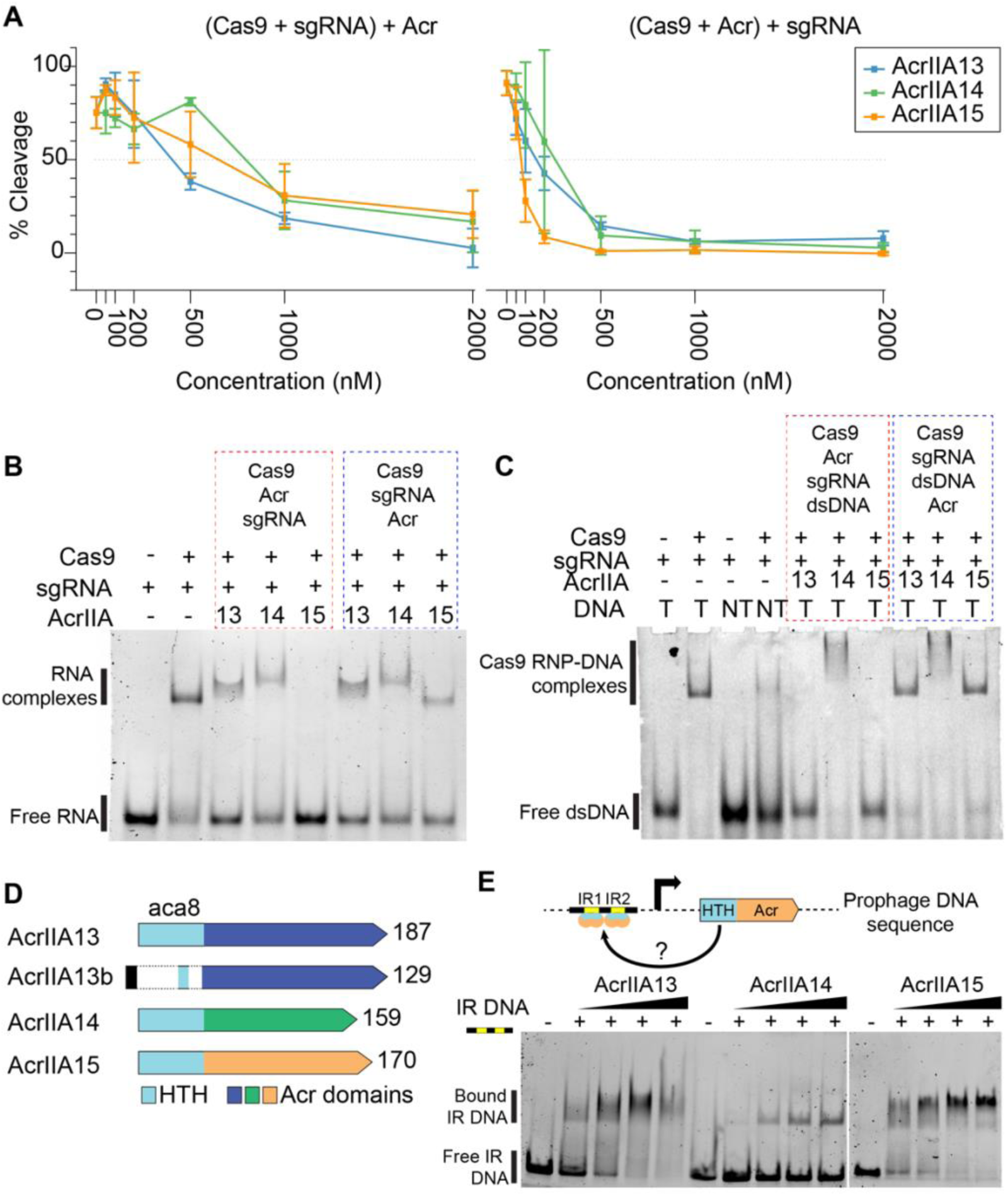
Three SauCas9 inhibitors with distinct features and activities. (A) Plot showing inhibition of SauCas9 cleavage activity by AcrIIA13-15 with % cleavage plotted on y-axis and Acr concentration (nM) on the x-axis. (Left) Plot corresponding to the cleavage assays performed by first complexing Cas9 and sgRNA followed by addition of Acr and DNA. (Right) Plot corresponding to the cleavage assays performed by first complexing Cas9 with Acrs followed by the addition of sgRNA and target dsDNA. The plotted data is the average % cleavage activity from 3 independent replicates. Error bars represent average +- sd. (B) 6% polyacrylamide gel showing formation of Cas9-sgRNA RNP in the presence and absence of different Acrs. Lanes boxed in red represent reactions where Acrs were added before the addition of sgRNA. Lanes boxed in blue represent reactions where Acrs were added after the addition of sgRNA. Order of the addition of different reaction components are shown above the boxes. (C) 6% polyacrylamide gel showing binding of Cas9-sgRNA RNP to either target DNA (T) or non-targeting DNA (NT) in the presence and absence of different Acrs. Lanes boxed in red represent reactions where Acrs were added to Cas9 before the addition of sgRNA and dsDNA. Lanes boxed in blue represent reactions where Acrs were added after the addition of dsDNA to Cas9-sgRNA RNP. Order of the addition of different reaction components are shown above the boxes. (D) Schematic showing the gene structure of AcrIIA13-15. The gene section highlighted in blue is highly conserved across the different Acrs and contains the conserved Helix-turn-helix (HTH) motif. The C-terminal region shows high variability across the three Acrs. (E) Schematic showing our hypothesis that the Acrs containing HTH motif binds to the promoter proximal sequence spanning inverted repeats -IR1 and IR2. 6% polyacrylamide gel showing binding of individually purified Acr proteins to AcrIIA15 promoter proximal sequence containing inverted repeats. The reaction components are indicated above the gel.

We then performed RNA electrophoretic mobility shift assays (EMSA) to determine if the Acrs were able to prevent, or reverse, Cas9-sgRNA RNP formation. While AcrIIA13 and AcrIIA14 did not prevent RNP formation, the addition of AcrIIA15 before the sgRNA during complex formation eliminated the presence of the slow-migrating species corresponding to Cas9 RNP formation that is visible when the Acr is added after RNP complex formation (Fig. 3B). The lack of a mobility-shifted species when AcrIIA15 is added before sgRNA suggests that AcrIIA15 binds directly to Cas9 to prevent RNP formation rather than causing RNP disassembly. Next, we performed EMSAs to evaluate the ability of the Acrs to prevent binding of the Cas9-sgRNA RNP to its target dsDNA. We found that both AcrIIA13 and AcrIIA15 inhibited dsDNA binding of Cas9, but only when added prior to the addition of target dsDNA. Interestingly, although AcrIIA14 completely inhibited Cas9 cleavage activity (Fig. 3A), it did not prevent Cas9 RNP from binding to its target. However, a super-shifted species was observed after the addition of AcrIIA14 and either before or after addition of dsDNA. This suggests that AcrIIA14 may act by binding the SauCas9 active site without impacting DNA binding, similar to the behavior of AcrIIC1 (31). AcrIIA14 might also trigger complex dimerization to form an inactive conformation, similar to the mechanisms of AcrIIC3 (31, 32) or AcrVA4 (33) (Fig. 3C). Altogether, these biochemical data suggest that each of the newly discovered Acrs inhibits SauCas9-mediated DNA cleavage by a different method.

### AcrIIA13-15 share a N-terminal domain that binds an inverted repeat

To determine the level of sequence similarity between AcrIIA13-15, we performed a multiple sequence alignment with MUSCLE and observed a shared N-terminal domain across all three Acrs, which was absent in AcrIIA13b, a truncated homolog of AcrIIA13 found during the guilt-by-association search (Fig. 3D). The remaining C-terminus, however, was not closely related between AcrIIA13-15 (Fig. S5A). Given the apparently distinct mechanisms of inhibition for AcrIIA13-15 respectively, we suspected that the residues downstream of the shared N-terminus were responsible for the Acr activity. To test this, we cloned, expressed and purified truncated versions of all three Acrs with the shared N-termini removed (Fig. S5A). In DNA cleavage assays with the purified C-terminal AcrIIA13-15 proteins, all the Acrs showed complete inhibition of Cas9 cleavage activity at ∼5-fold molar excess whether added before or after Cas9-sgRNA RNP formation (Fig. S5B). We also observe strong inhibition of the C-terminal portion of AcrIIA13 in the TXTL sfGFP expression assay, while no inhibition was detected using the N-terminal domain alone (Fig. S5C). Overall, these data support the conclusion that the variable C-terminal domains of AcrIIA13-15 constitute the Acr activity of each protein. We also tested the truncated AcrIIA13-15 proteins for inhibition of SpyCas9 in an RNA-guided DNA cleavage assay, but observed no significant reduction in DNA cutting (Fig. S5D).

### The shared Acr N-terminal domain binds an inverted-repeat DNA sequence

On further sequence analysis of the homologous N-terminal section of AcrIIA13-15, we found they contain a conserved helix-turn-helix (HTH) motif (Fig. 3D and Fig. S5A). These HTH domains have been found in aca genes and were recently shown to repress their own transcription along with other Acrs in the same operon (16, 34). These aca genes encode proteins that bind to promoter-proximal sequences containing two inverted repeats to block polymerase access. We observe inverted repeats (IR) proximal to the promoters driving transcription of the operons containing AcrIIA13-15 (Fig. S5A). To determine if AcrIIA13-15 are capable of binding to the promoter proximal IR sequences, we performed DNA EMSAs, combining each purified Acr protein with a FAM-labelled dsDNA sequence spanning the IRs in the promoter-proximal region of AcrIIA15 (Fig. S5B). We found that AcrIIA13 and AcrIIA15 strongly interact with the promoter-proximal IR sequence, while AcrIIA14 only weakly interacted with it (Fig. 3E). Mutating the IR sequence of the dsDNA substrate completely abolished the Acr-IR DNA interaction. To further assess if this interaction was mediated by the HTH motif, we repeated the DNA EMSA using the C-terminal AcrIIA15, where we failed to detect any interaction with the dsDNA IR-DNA substrate (Fig. S6C). Together, these data suggest that, similar to AcrIIA1 (16), AcrIIA13-15 also are fusion proteins that link an anti-SauCas9 Acr with a DNA binding domain that is likely to regulate the expression of its own operon.

### Inhibition of SauCas9-mediated genome editing in human cells

To assess the ability of AcrIIA13-15 to inhibit genome editing in the context of human cells, we designed a lentiviral validation system to express Cas9, an sgRNA, and an Acr protein from three different, stably integrated constructs (Fig. 4A). Using HEK-RT1 genome editing reporter cells (35, 36) - a monoclonal HEK293T-based cell line expressing a doxycycline-inducible GFP reporter - lentiviral constructs encoding the AcrIIA proteins or mTagBFP2, and SauCas9 or SpyCas9, were sequentially transduced and stable integrants selected. The resulting cell lines were then further transduced with a vector expressing a GFP-targeting or negative control sgRNA and an associated mCherry fluorescence marker. In this system, we observed a baseline of ∼75% GFP-positive cells after induction in presence of a negative control sgRNA, which was reduced to near 0% by an on-target sgRNA disrupting GFP expression (Fig. S7).

**Figure 4.**
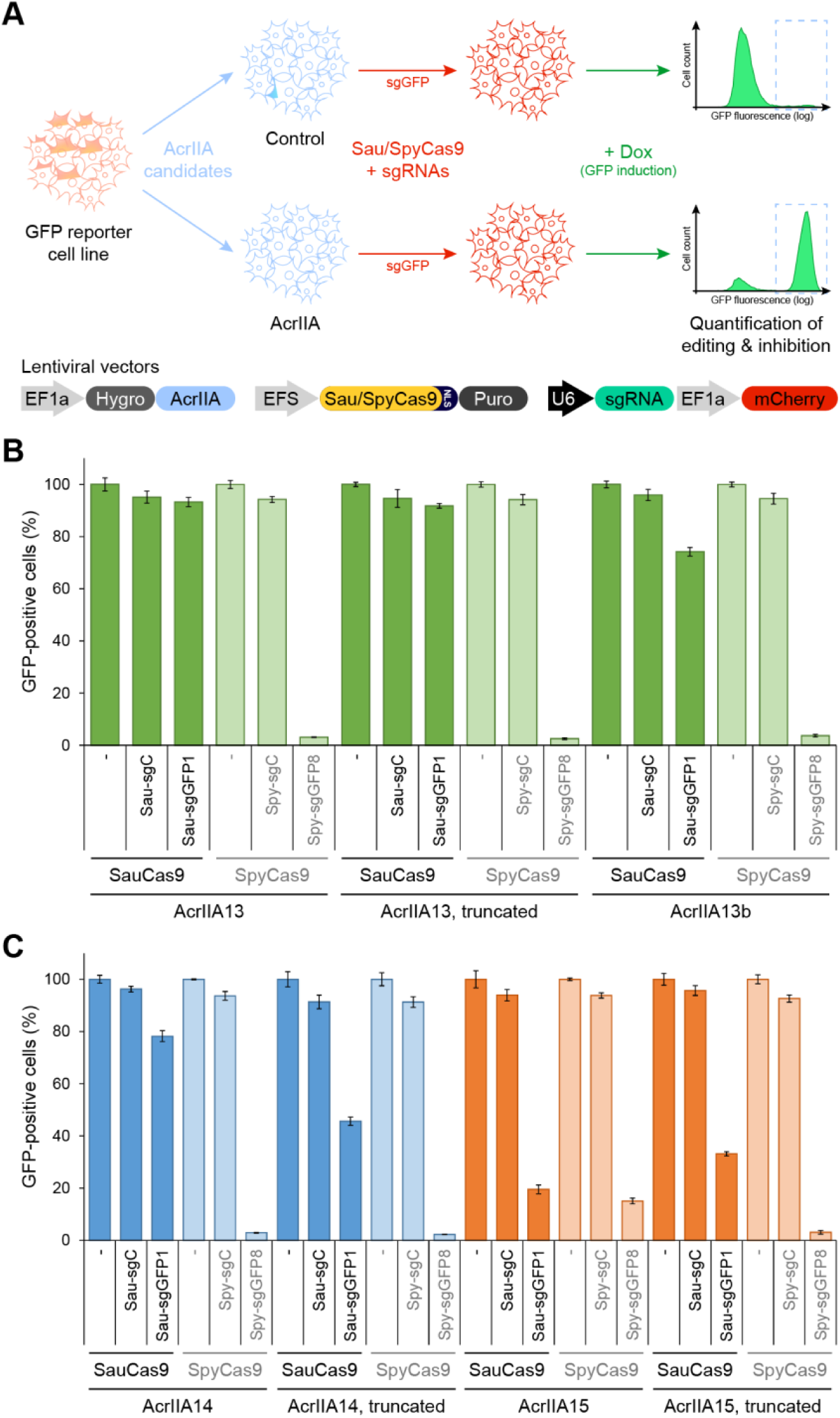
AcrIIA13 is a potent and selective inhibitor of SauCas9 in mammalian cells. (A) Schematic of a mammalian validation platform to express Cas9, sgRNAs, and an Acr protein from different stably integrated lentiviral vectors in human cells, to quantify the CRISPR-Cas genome editing inhibitory potential of the Acr candidates by flow cytometry. HEK-RT1 cells, a human HEK293T-based genome editing reporter cell line with a doxycycline-inducible GFP, were sequentially transduced with lentiviral vectors expressing Acr candidates, SauCas9 or SpyCas9, and guide RNAs targeting GFP or a negative control. Three days post-transduction of the guide RNAs, GFP expression was induced by treatment with doxycycline for 24 hours. Genome editing inhibition efficiency for each Acr candidate was measured as the increase in percentage GFP-positive cells expressing on-target guide RNAs (sgGFP, mCherry-positive), compared to controls. (B) Quantification of SauCas9 and SpyCas9 inhibition efficiency of AcrIIA13, an N-terminal truncation of AcrIIA13, and an AcrIIA13 homolog (AcrIIA13b) in human cells. Error bars represent the standard deviation of triplicates. sgC, sgGFP1/sgGFP8: negative control and GFP-targeting guide RNAs for the respective Cas9s. The data shown for AcrIIA13 are partially quantification of raw flow cytometry data shown in Fig. S9. (C) Assessment of SauCas9 and SpyCas9 inhibition by AcrIIA14, an N-terminal truncation of AcrIIA14, AcrIIA15, and an N-terminal truncation of AcrIIA15 in human cells. The assay was run as described above.

We next tested three known Acrs for their ability to inhibit SauCas9-mediated genome editing in mammalian cells, including the robust SpyCas9 inhibitor AcrIIA4 (10), the broad-spectrum type II-A Cas9 inhibitor AcrIIA5 (12) and the species-specific *Streptococcus thermophilus* Cas9 inhibitor AcrIIA6 (12). While AcrIIA4 prevented genome editing by SpyCas9 as expected (10), we found that it had no effect against SauCas9 (Fig. S8). Conversely, AcrIIA5 displayed strong inhibitory activity towards both SauCas9 and SpyCas9, while AcrIIA6 did not show any inhibition of either of the two Cas9s.

Similar genome editing inhibition experiments using AcrIIA13-15 showed that AcrIIA13 is a potent inhibitor of SauCas9, with near complete inhibition of GFP gene editing activity (Fig. 4B and Fig. S9). However, in contrast to AcrIIA5, AcrIIA13 did not inhibit SpyCas9, making it the first Acr selective for SauCas9 to our knowledge, and further expanding the CRISPR-Cas modulation toolkit. Interestingly, the truncated form of AcrIIA13, which is substantially smaller, was equally effective at inhibiting SauCas9-mediated genome editing in human cells. The AcrIIA13b homolog that lacks the N-terminal DNA-binding domain showed a comparable trend of SauCas9 selective inhibition. Full-length AcrIIA14 also proved to be a potent inhibitor of SauCas9, although less robust than AcrIIA13, and removing its N-terminal domain had a larger impact on its inhibitory efficacy (Fig. 4C). SpyCas9 was not affected by AcrIIA14. Finally, only AcrIIA15 demonstrated any activity toward SpyCas9-mediated genome editing, but its inhibition was minimal (Fig. 4C), showing that AcrIIA13-15 are potent and selective inhibitors for SauCas9.

## Discussion

The potential role of anti-CRISPRs in regulating CRISPR-Cas gene editing in therapeutic applications has helped spur the discovery of over 40 different families of Acrs (3). However, creating ‘designer’ inhibitors for specific gene editors and editing scenarios is very challenging as Acr proteins are small and lack any sequence and structural conservation to use for direct identification of desired modalities. To overcome these problems, approaches such as screening self-targeting genomes (7, 10, 19) or metagenomic DNA (13, 14) attempt to cast a wide net in the genomic space to identify novel Acr activities. A more direct method is to use a ‘guilt-by-association’ search approach, which leverages the repeated observations that Acrs tend to co-exist in similar operons and that the genomic neighborhood of several Acrs contain anti-CRISPR associated (*aca*) genes that are closely coupled with Acrs, and widespread in bacteria and MGEs.

In this work, we sought to identify potent inhibitors of SauCas9, a common SpyCas9 alternative. Using a combination of the methods described above, we identified three novel families of Acrs (AcrIIA13-15) that are potent inhibitors of SauCas9. Using biochemical approaches, we find that all three Acrs possess different mechanisms of Cas9 inhibition. AcrIIA13 prevents SauCas9-mediated dsDNA cleavage activity by blocking DNA binding, while AcrIIA15 binds to SauCas9 to block RNP assembly (Figs. 3B, 3C). Our data also show that AcrIIA14 does not interfere with SauCas9 binding to either RNA or dsDNA, but inhibits dsDNA cleavage activity at a potency comparable to AcrIIA13 and AcrIIA15. It is possible that AcrIIA14 either causes the SauCas9-sgRNA RNP to form higher order structures like AcrVA4 (33) or AcrIIC3 (31, 32), or interacts with RNP to prevent cleavage but not binding like AcrIIC1 (31). We also observed potent and SauCas9-specific inhibition by AcrIIA13 and AcrIIA14 in mammalian cells (Fig, 4). AcrIIA15 was able to inhibit both SpyCas9 and SauCas9 in human cells, but to a much lesser extent. Of the three Acrs, AcrIIA13 was the most potent inhibitor of SauCas9 in human cells.

In comparing the protein sequences between AcrIIA13-15, we observe that the N-termini of the Acrs possessed a highly conserved HTH motif (Figs. 3D and S4A). This motif is a prominent feature of *aca* genes, which were recently shown to negatively regulate the transcription of their own operon (16, 34). The self-regulating activity of aca proteins was shown to be mediated by a direct interaction between the HTH motif and promoter proximal sequences containing two inverted repeats (IR). Examining the promoter regions of AcrIIA13-15, we found that all of them also contain similar promoter-proximal IR sequences (Fig. S6A), and that all three Acrs are able to bind to the IR sequences specifically and only when the N-terminal domain is present (Fig. 3E and S6C). The DNA binding activity of the N-terminal region also has little to no effect on Acr potency when removed from the protein in *in vitro* cleavage assays (Fig. S5B-D) and no or only a minor effect in human cells depending on the Acr (Fig. 4B,C). AcrIIA13 in particular is unaffected by the removal of its N-terminus, and represents a highly potent, SauCas9-specific inhibitor for use in mammalian contexts for regulating Cas9 activity.

Based on our data, the AcrIIA13-15 proteins we identified are likely fusions between three unique Acr proteins and a shared *aca* gene, which we name *aca8*. It is unclear how aca8 became fused with different Acr proteins, or what benefit a single protein would have over the split alternative. Regardless, the new aca8 gene provides another seed to search for further discoveries using the guilt-by-association approach. With the addition of yet more proteins, we increase the chances of finding new activities that are particularly attractive to biotechnology and therapeutic applications, such as catalytic Acrs that would have low delivery requirements for potent inhibition (37, 38). We believe that the search for more Acrs with potent and novel mechanisms of inhibition will continue to rapidly add new dimensions of control into the CRISPR toolbox.

## Materials and Methods

### Self-targeting strain selection

The bioinformatics search for self-targeting *Staphylococcus* strains was carried out in a similar manner as described in Watters, et al. (19). Briefly, the Self-Targeting Spacer Searcher (STSS) (https://github.com/kew222/Self-Targeting-Spacer-Searcher) was used the NCBI database for all assemblies returned using a search of “Staphylococcus[organism] NOT phage NOT virus”. Each assembly was checked for instances of self-targeting, which were reported along with details about the instance including: target site mutations, up/downstream target sequences, Cas genes found near the array of the self-targeting spacer, predicted CRISPR subtype based on repeats/Cas proteins, repeat sequences and conservation, etc. Systems that were output as ambiguous type II CRISPR systems were manually checked for: 1) repeat sequence similarity to other known loci, 2) presence of defining Cas proteins (i.e. Csn2 for II-A), and 3) alignment of Cas9 of the system in question to a set of type II-A, II-B, and II-C Cas9s to determine similarity. The final set of self-targeting instances found in *Staphylococcus* genomes on NCBI can be found in Table S1. Strains were chosen for further study based on similarity of the downstream target site to the SauCas9 PAM (3′-NNGRR(T)) and similarity of the self-targeting system’s Cas9 to SauCas9 by blastp.

### Guilt-by-association search

GF genes 1 and 2 (AcrIIA13) were used to query the NCBI protein database with blastp to identify homologs. Each homolog found was then manually surveyed for neighboring genes that were not found within the genome fragment amplicons tested with the TXTL sfGFP assay. This approach was repeated using any neighboring proteins to further locate their neighbors. Only alignments with an E-value < 0.001 were considered. The search was ended after 10 candidate proteins were identified.

### gDNA extraction

To extract gDNA, 4 mL of a culture containing *Staphylococcus haemolyticus* (strains W_75 and W_139) or *Staphylococcus schleiferi* (strains 5909-02 and 2713-03) cells were grown overnight in Tryptic Soy Broth. The cultures were then pelleted and resuspended in 500 μL of PBS. Cells were then treated with lysostaphin for 1 hr at 37°C. gDNA was then extracted using the Monarch gDNA Purification Kit from NEB following the manufacturer’s instructions.

### Template preparation for TXTL

Each TXTL reaction contained three template DNA sources: a sfGFP constitutive reporter plasmid (pUC), a plasmid expressing a sgRNA targeting GFP (pUC) using the SauCas9 sgRNA sequence from Strutt, et al. (27), an either a plasmid expressing SauCas9 under an arabinose promoter (p15A) or a genomic amplicon containing the Cas9 from *S. haemolyticus* or *S. schleiferi*. Reactions may also contain an Acr or Acr candidate expressed from a Tet-repressible promoter or a genomic fragment amplicon with the naturally occurring sequence. The plasmid preparation for the reporter, sgRNA plasmid, and SauCas9 were as described in Watters, et al. (19). To prepare the Cas9 genomic amplicons, PCR of purified gDNA with Q5 polymerase (New England Biolabs) was performed with primers that spanned all of the Cas genes (Table S3). Similarly, genomic fragment amplicons were prepared with PCR using the gDNA and primers listed in Table S3. Acr candidates that were to be tested individually were cloned into a plasmid for use in TXTL.

To prepare the plasmids for TXTL, a 20 mL culture of *E. coli* containing one of the plasmids was grown to high density, then isolated across five preparations using the Monarch Plasmid Miniprep Kit (New England Biolabs), eluting in a total of 200 μL nuclease-free H_2_O. 200 μL of AMPure XP beads (Beckman Coulter) were then added to each combined miniprep and purified according to the manufacturer’s instructions, eluting in a final volume of 20 μL in nuclease-free H_2_O.

All anti-CRISPR candidate amplicons and subfragments were prepared using 100 μL PCRs with Q5 or Phusion polymerase (New England Biolabs), under various conditions to yield a clear single band on an agarose gel such that the correct fragment length was greater than 95% of the fluorescence intensity of the lane on the gel. 100 μL of AMPure XP beads (Beckman Coulter) were then added to each reaction and purified according to the manufacturer’s instructions, eluting in a final volume of 10 μL in nuclease-free H_2_O. The PCRs and purification of the Cas9 genomic amplicons was scaled up 5-fold. A listing of the plasmids used in the TXTL experiments can be found in Table S5.

### TXTL reactions

The reactions were carried out as described in Watters, et al. (19). Briefly, each reaction was mixed from 9 μL of TXTL master mix, 0.125 nM of each reporter plasmid, 1 nM of Cas9 amplicon or plasmid, 2 nM of sgRNA plasmid, 1 nM of genomic fragment amplicon or Acr candidate plasmid, 1 μM of IPTG, 0.5 μM of anhydrotetracycline, 0.1% arabinose, and 2 μM of annealed oligos containing six χ sites (39).

The reactions were run at 29 °C in a TECAN Infinite Pro F200, measuring GFP (λex: 485 nm, λem: 535 nm) fluorescence levels every three minutes for 10 hours. To plot kinetic data, the minimum measured fluorescence intensity was subtracted from each point on the curve (to compensate for early variations due to condensation on the sealing film), then the overall curve was normalized by the fluorescence level measured for the non-targeting negative control after 10 hours of reporter expression.

### Protein purification

DNA sequences corresponding to SauCas9 and variants of AcrIIA13-15 were cloned downstream of 10xHis tag, MBP tag and a TEV cleavage site. Each of these constructs were grown in Rosetta2 cells overnight in Lysogeny Broth and were subcultured in 750 mL of Terrific Broth. After the cultures reached an OD_600_ of 0.6-0.8, proteins were induced by treatment with 0.5 mM IPTG for ∼20 hrs at 16°C. Cells were then resuspended in lysis buffer containing 50 mM HEPES pH 7.0, 500 mM NaCl, 1 mM TCEP and 5% glycerol and Roche cOmplete protease inhibitor cocktail and lysed by sonication, followed by purification using Ni-NTA Superflow resin (Qiagen). The eluted proteins were then cleaved with TEV protease overnight at 4°C while performing buffer exchange in buffer containing 250 mM NaCl. The cleaved proteins were further purified using Ion exchange chromatography with a linear NaCl gradient using Heparin HiTrap column (GE) or HiTrap Q column (GE). The protein samples were then concentrated and subjected to size exclusion chromatography with Superdex 200 size exclusion column (GE). The proteins were eluted in final storage buffer containing 20 mM HEPES pH 7.0, 200 mM KCl, 1 mM TCEP and 5% glycerol. Plasmids for expressing the Acrs are available on Addgene with ##XXX, XXX, XXX.

### In vitro cleavage assays

All dsDNA cleavage assays were performed in a 1X Cleavage Buffer containing 20 mM HEPES-HCl, pH 7.5, 150 mM KCl, 10 mM MgCl_2_, 0.5 mM TCEP. sgRNA sequences synthesized by IDT were first refolded in 1X cleavage buffer by heating at 70°C for 5 min then cooling to room temperature. For all the cleavage assays involving SauCas9, reaction components were added in the following two ways: (1) SauCas9 and Acrs were complexed together, followed by sequential addition of sgRNA and dsDNA. (2) SauCas9 and sgRNA were complexed together, followed by addition of Acrs and dsDNA target. Each step of sequential additions in both reactions were preceded by incubation at 37°C for 10 min. All cleavage reactions were then quenched on completion with 2 µL of 6X Quenching Buffer containing 30% glycerol, 1.2% SDS, 250 mM EDTA. Cleavage products were read using a 1% agarose gel prestained with SYBRGold. All DNA/RNA sequences used for biochemical assays can be found in Table S6.

### Electrophoretic mobility shift assays

RNA EMSAs were performed by incubating SauCas9 and sgRNA in 1X cleavage buffer lacking MgCl_2_ in the presence or absence of 10-fold molar excess of different Acrs relative to SauCas9. The reactions were then incubated at 37°C for 10 mins and run on 6% TBE polyacrylamide gels for 1 hr at 50V. The gels were then visualized using SYBRGold stain. DNA EMSAs to evaluate binding of SauCas9 RNP to dsDNA target were performed by incubating the reaction components described in their respective figures in 1X cleavage buffer lacking MgCl_2_ at 37°C for 10 min. The dsDNA substrates used for this assay were synthesized as 5’ FAM labelled 34 bp DNA oligos that were annealed prior to their addition to the EMSA reaction. The reactions were run on 6% TBE polyacrylamide gels and visualized on a Typhoon imager (GE). DNA EMSAs for the binding of AcrIIA13-15 to IR DNA were performed by incubating the components in 1X cleavage buffer lacking MgCl_2_ at 37°C for 10 min. The completed reactions were then run on 6% TBE polyacrylamide gels for 1 hr at 50V and visualized with SYBRGold stain.

### Lentiviral vectors

The lentiviral vector pCF525-mTagBFP2, expressing an EF1a-driven polycistronic construct containing a hygromycin B resistance marker, P2A ribosomal skipping element, and an mTagBFP2 fluorescence marker had been cloned before (19). For stable lentiviral expression of AcrIIA candidates, mTagBFP2 was replaced by the gene-of-interest open reading frame (ORF) using custom oligonucleotide gBlocks (IDT), Gibson assembly reagents (NEB), and standard molecular cloning techniques. The SpyCas9 guide RNA-only lentiviral vector pCF820, encoding a SpyCas9 U6-sgRNA cassette and an EF1a driven, human codon optimized mCherry2 marker (40), was based on the pCF221 vector (35). To make the backbone more efficient and increase viral titers, the f1 bacteriophage origin of replication and bleomycin resistance marker were removed, as has previously been done for pCF525 (19). The SauCas9 guide RNA-only lentiviral vector pCF824, encoding an SauCas9 U6-sgRNA cassette and an EF1a driven, human codon optimized mCherry2 marker, was based on the pCF820 vector. The SauCas9 guide RNA scaffold was designed as previously reported (41), synthesized as custom oligonucleotides (IDT), and assembled using Gibson techniques and standard molecular cloning methods. A lentiviral vector referred to as pCF823, expressing an EFS driven SpyCas9-P2A-PuroR cassette, was based on the vector pCF204 (35) by removing the f1 bacteriophage origin of replication and bleomycin resistance marker to increase viral titers. A lentiviral vector referred to as pCF825, expressing an EFS driven SauCas9-P2A-PuroR cassette, was based on the vector pCF823 by replacing the SpyCas9 ORF with the SauCas9 ORF from pX601 (Addgene #61591) (25). Vector sequences are provided in a separate file (Table S7).

### Design of sgRNAs

The following protospacer sequences were used for SpyCas9 sgRNAs: sgC (GGAGACGGAGGACGACGAACGTCTCT), sgGFP8 (GCAGGGTCAGCTTGCCGTAGG). The following sequences were used for SauCas9 sgRNAs: sgC (GGAGACGGAGGACGACGAACGTCTCT), sgGFP1 (GCAAGGGCGAGGAGCTGTTCAC).

### Mammalian cell culture

All mammalian cell cultures were maintained in a 37°C incubator at 5% CO_2_. HEK293T (293FT; Thermo Fisher Scientific) human kidney cells and derivatives thereof were grown in Dulbecco’s Modified Eagle Medium (DMEM; Corning Cellgro, #10-013-CV) supplemented with 10% fetal bovine serum (FBS; Seradigm #1500-500), and 100 Units/ml penicillin and 100 μg/ml streptomycin (100-Pen-Strep; Gibco #15140-122). HEK293T and HEK-RT1 cells were tested for absence of mycoplasma contamination (UC Berkeley Cell Culture facility) by fluorescence microscopy of methanol fixed and Hoechst 33258 (Polysciences #09460) stained samples.

### Lentiviral transduction

Lentiviral particles were produced in HEK293T cells using polyethylenimine (PEI; Polysciences #23966) based transfection of plasmids, as previously described (35). In brief, lentiviral vectors were co-transfected with the lentiviral packaging plasmid psPAX2 (Addgene #12260) and the VSV-G envelope plasmid pMD2.G (Addgene #12259). Transfection reactions were assembled in reduced serum media (Opti-MEM; Gibco #31985-070). For lentiviral particle production on 6-well plates, 1 µg lentiviral vector, 0.5 µg psPAX2 and 0.25 µg pMD2.G were mixed in 0.4 ml Opti-MEM, followed by addition of 5.25 µg PEI. After 20-30 min incubation at room temperature, the transfection reactions were dispersed over the HEK293T cells. Media was changed 12 h post-transfection, and virus harvested at 36-48 h post-transfection. Viral supernatants were filtered using 0.45 µm polyethersulfone (PES) membrane filters, diluted in cell culture media if appropriate, and added to target cells. Polybrene (5 µg/ml; Sigma-Aldrich) was supplemented to enhance transduction efficiency, if necessary.

### Gene editing inhibition assays in human cells

Candidate anti-CRISPR (Acr) proteins were tested in a HEK293T-based monoclonal reporter cell line called HEK-RT1 (35, 36), which features a doxycycline-inducible GFP reporter (42) that can be edited before doxycycline-based induction of its expression. To test the effect of genomic integration and expression of AcrIIA candidate proteins in mammalian cells, HEK-RT1 cells were stably transduced with the lentiviral vector pCF525 (19) encoding the various candidate proteins or mTagBFP2, and selected on hygromycin B as previously described (19). The resulting HEK-RT1-AcrIIA candidates and HEK-RT1-mTagBFP2 genome protection and editing reporter cell lines were then used to quantify genome editing inhibition by flow cytometry. For this, all reporter cell lines were stably transduced with the lentiviral vector pCF823-SpyCas9 (expressing SpyCas9-P2A-PuroR from an EFS promoter) or pCF825-SauCas9 (expressing SauCas9-P2A-PuroR from an EFS promoter) and selected on puromycin (1.0 µg/ml). AcrIIA/mTagBFP2 and SpyCas9/SauCas9 expressing HEK-RT1 reporter cell lines were then stably transduced with lentiviral vectors pCF820-Spy-sgGFP8/sgC (U6 promoter driven expression of SpyCas9-specific guide RNAs targeting GFP or a non-targeting recipient control, along with an EF1a driven humanized mCherry2 marker) or pCF824-Sau-sgGFP1/sgC (U6 promoter driven expression of SauCas9-specific guide RNAs targeting GFP or a non-targeting recipient control, along with an EF1a driven humanized mCherry2 marker). At day three post-transduction of the guide RNA vectors, anti-CRISPR (AcrIIA candidates, mTagBFP2) + Cas9 (SpyCas9, SauCas9) + guide RNA (Spy-sgC/sgGFP8, Sau-sgC/sgGFP1) expressing HEK-RT1 reporter cells were treated with doxycycline (1 µg/ml) for GFP reporter induction. At 24 h post-induction, transgenic GFP genome editing efficiency and inhibition thereof were quantified by flow cytometry (Attune NxT, Thermo Fisher Scientific). For guide RNA expressing samples, quantification was gated on the mCherry-positive (lentiviral guide RNA transduced) population. Data was normalized on samples without guide RNA expression treated +/- doxycycline (1 µg/ml).

## Acknowledgements

We thank Hua B. Bai for cloning plasmids used in the TXTL assay, Shawn M. Ren for help with preliminary cell line construction, Fuguo Jiang for a gift of AcrIIA4 protein, Gavin Knott for a gift of AcrVA4 protein, the Bondy-Denomy Lab for sharing their lab space, and the Doudna Lab for thoughtful comments and feedback. The authors also thank Kathleen Boyajian and Shelley Rankin at the University of Pennsylvania for the generous gift of the *S. schleiferi* strains, and Jorunn Pauline Cavanagh from the University of Norway, Tromsø for the generous gift of the *S. haemolyticus* strains. The authors acknowledge financial support from the Defense Advanced Research Projects Agency (DARPA) (award HR0011-17-2-0043 to J.A.D.), the Paul G. Allen Frontiers Group and the National Science Foundation (MCB-1244557 to J.A.D.). C.F. is supported by a US National Institutes of Health K99/R00 Pathway to Independence Award (K99GM118909, R00GM118909) from the National Institute of General Medical Sciences (NIGMS). J.A.D. is an investigator of the Howard Hughes Medical Institute (HHMI), and this study was supported in part by HHMI; J.A.D is also a Paul Allen Distinguished Investigator.

## Author contributions

Conceptualization, K.E.W.; Methodology, K.E.W., H.S., C.F.; Software, K.E.W.; Investigation, K.E.W., H.S., C.F., R.J.L., B.M.; Data Curation, K.E.W., H.S., C.F.; Writing, K.E.W., H.S., C.F., J.A.D.; Funding Acquisition, K.E.W., C.F., J.A.D.

## References

1. Knott GJ, Doudna JA (2018) CRISPR-Cas guides the future of genetic engineering. Science 361(6405):866–869.

2. Zhang F, Song G, Tian Y (2019) Anti-CRISPRs: The natural inhibitors for CRISPR-Cas systems. Anim Models Exp Med 482(7385):331–7.

3. Hwang S, Maxwell KL (2019) Meet the Anti-CRISPRs: Widespread Protein Inhibitors of CRISPR-Cas Systems. The CRISPR Journal 2(1):23–30.

4. Bondy-Denomy J, Pawluk A, Maxwell KL, Davidson AR (2013) Bacteriophage genes that inactivate the CRISPR/Cas bacterial immune system. Nature 493(7432):429–432.

5. Pawluk A, Bondy-Denomy J, Cheung VHW, Maxwell KL, Davidson AR (2014) A new group of phage anti-CRISPR genes inhibits the type I-E CRISPR-Cas system of Pseudomonas aeruginosa. MBio 5(2):e00896.

6. Pawluk A, et al. (2016) Inactivation of CRISPR-Cas systems by anti-CRISPR proteins in diverse bacterial species. Nature Microbiology 1(8):16085.

7. Marino ND, et al. (2018) Discovery of widespread Type I and Type V CRISPR-Cas inhibitors. Science:eaau5174.

8. He F, et al. (2018) Anti-CRISPR proteins encoded by archaeal lytic viruses inhibit subtype I-D immunity. Nature Microbiology:1–11.

9. Pawluk A, et al. (2016) Naturally Occurring Off-Switches for CRISPR-Cas9. Cell 167(7):1829–1834.e9.

10. Rauch BJ, et al. (2017) Inhibition of CRISPR-Cas9 with Bacteriophage Proteins. Cell 168(1-2):150–158.e10.

11. Hynes AP, et al. (2017) An anti-CRISPR from a virulent streptococcal phage inhibits Streptococcus pyogenes Cas9. Nature Microbiology 2(10):1374–1380.

12. Hynes AP, et al. (2018) Widespread anti-CRISPR proteins in virulent bacteriophages inhibit a range of Cas9 proteins. Nature Communications 9(1):2919–10.

13. Uribe RV, et al. (2019) Discovery and Characterization of Cas9 Inhibitors Disseminated across Seven Bacterial Phyla. Cell Host and Microbe:1–20.

14. Forsberg KJ, et al. (2019) Functional metagenomics-guided discovery of potent Cas9 inhibitors in the human microbiome. elife 8:1709.

15. Lee J, et al. (2018) Potent Cas9 Inhibition in Bacterial and Human Cells by AcrIIC4 and AcrIIC5 Anti-CRISPR Proteins. MBio 9(6):e02321–18.

16. Osuna BA, Karambelkar S, Mahendra C, bioRxiv KC, 2019 Listeria phages induce Cas9 degradation to protect lysogenic genomes. biorxivorg. doi:10.1101/787200.

17. Bhoobalan-Chitty Y, Johansen TB, Di Cianni N, Peng X (2019) Inhibition of Type III CRISPR-Cas Immunity by an Archaeal Virus-Encoded Anti-CRISPR Protein. Cell 179(2):448–458.e11.

18. Athukoralage JS, Graham S, Grüschow S, Rouillon C, White MF (2019) A Type III CRISPR Ancillary Ribonuclease Degrades Its Cyclic Oligoadenylate Activator. Journal of Molecular Biology 431(15):2894–2899.

19. Watters KE, Fellmann C, Bai HB, Ren SM, Doudna JA (2018) Systematic discovery of natural CRISPR-Cas12a inhibitors. Science 9:eaau5138.

20. Rauch BJ, et al. (2017) Inhibition of CRISPR-Cas9 with Bacteriophage Proteins. Cell 168(1-2):150–158.e10.

21. Heussler GE, O’Toole GA (2016) Friendly Fire: Biological Functions and Consequences of Chromosomal Targeting by CRISPR-Cas Systems. Journal of Bacteriology 198(10):1481–1486.

22. Bondy-Denomy J, et al. (2018) A Unified Resource for Tracking Anti-CRISPR Names. The CRISPR Journal 1(5):304–305.

23. Shin J, et al. (2017) Disabling Cas9 by an anti-CRISPR DNA mimic. Sci Adv 3(7):e1701620.

24. Wang Q, et al. (2018) Genome scale screening identification of SaCas9/gRNAs for targeting HIV-1 provirus and suppression of HIV-1 infection. Virus Research:1–36.

25. Ran FA, et al. (2015) In vivo genome editing using Staphylococcus aureus Cas9. Nature 520(7546):186–191.

26. Zhao X, Yu Z, Xu Z (2018) Study the Features of 57 Confirmed CRISPR Loci in 38 Strains of Staphylococcus aureus. Front Microbiol 9:280–14.

27. Strutt SC, Torrez RM, Kaya E, Negrete OA, Doudna JA (2018) RNA-dependent RNA targeting by CRISPR-Cas9. elife 7:e32724.

28. Marshall R, et al. (2018) Rapid and Scalable Characterization of CRISPR Technologies Using an E. coli Cell-Free Transcription-Translation System. Molecular Cell 69(1):146–157.e3.

29. Arndt D, et al. (2016) PHASTER: a better, faster version of the PHAST phage search tool. Nucleic Acids Research 44(W1):W16–21.

30. Hudson CM, Lau BY, Williams KP (2015) Islander: a database of precisely mapped genomic islands in tRNA and tmRNA genes. Nucleic Acids Research 43(Database issue):D48–53.

31. Harrington LB, et al. (2017) A Broad-Spectrum Inhibitor of CRISPR-Cas9. Cell 170(6):1224–1233.e15.

32. Zhu Y, et al. (2019) Diverse Mechanisms of CRISPR-Cas9 Inhibition by Type IIC Anti-CRISPR Proteins. Molecular Cell 74(2):296–309.e7.

33. Knott GJ, et al. (2019) Structural basis for AcrVA4 inhibition of specific CRISPR-Cas12a. elife 8:213.

34. Stanley SY, et al. (2019) Anti-CRISPR-Associated Proteins Are Crucial Repressors of Anti-CRISPR Transcription. Cell 178(6):1452–1464.e13.

35. Oakes BL, et al. (2019) CRISPR-Cas9 Circular Permutants as Programmable Scaffolds for Genome Modification. Cell 176(1-2):254–267.e16.

36. Park HM, et al. (2018) Extension of the crRNA enhances Cpf1 gene editing in vitro and in vivo. Nature Communications 9(1):3313–12.

37. Knott GJ, et al. (2019) Broad-spectrum enzymatic inhibition of CRISPR-Cas12a. Nature Structural &amp; Molecular Biology 26(4):1–11.

38. Dong L, et al. (2019) An anti-CRISPR protein disables type V Cas12a by acetylation. Nature Structural &amp; Molecular Biology 26(4):1–15.

39. Marshall R, Maxwell CS, Collins SP, Beisel CL, Noireaux V (2017) Short DNA containing χ sites enhances DNA stability and gene expression in E. coli cell-free transcription-translation systems. Biotechnol Bioeng 114(9):2137–2141.

40. Ai H-W, Shaner NC, Cheng Z, Tsien RY, Campbell RE (2007) Exploration of new chromophore structures leads to the identification of improved blue fluorescent proteins. Biochemistry 46(20):5904–5910.

41. Chen B, et al. (2016) Expanding the CRISPR imaging toolset with Staphylococcus aureus Cas9 for simultaneous imaging of multiple genomic loci. Nucleic Acids Research 44(8):e75.

42. Fellmann C, et al. (2013) An optimized microRNA backbone for effective single-copy RNAi. Cell Rep 5(6):1704–1713.

