## Supplemental Information for "Potent CRISPR-Cas9 inhibitors from *Staphylococcus* genomes"

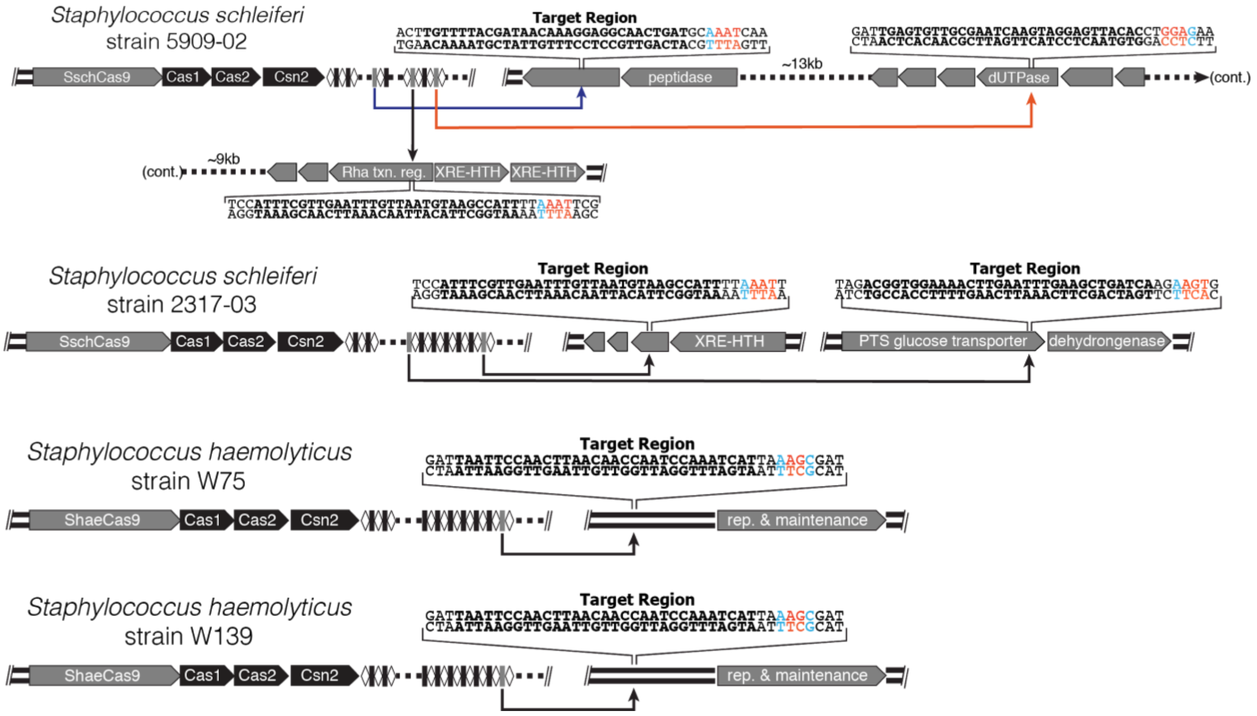


**Figure S1. Self-targeting spacers of *S. schleiferi* and *S. haemolyticus*.** The two *S. schleiferi* strains of interest contain self-targets that occur within a predicted prophage, with strain 5909-02 containing three self-targeting spacers and strain 2317-03 containing two. Both strains of *S. haemolyticus* selected for further study contained the same exact self-targeting spacer. The sequences downstream of the target are colored based on their similarity to the SauCas9 NNGRR(T) PAM where bases in red match the PAM motif and blue bases do not.

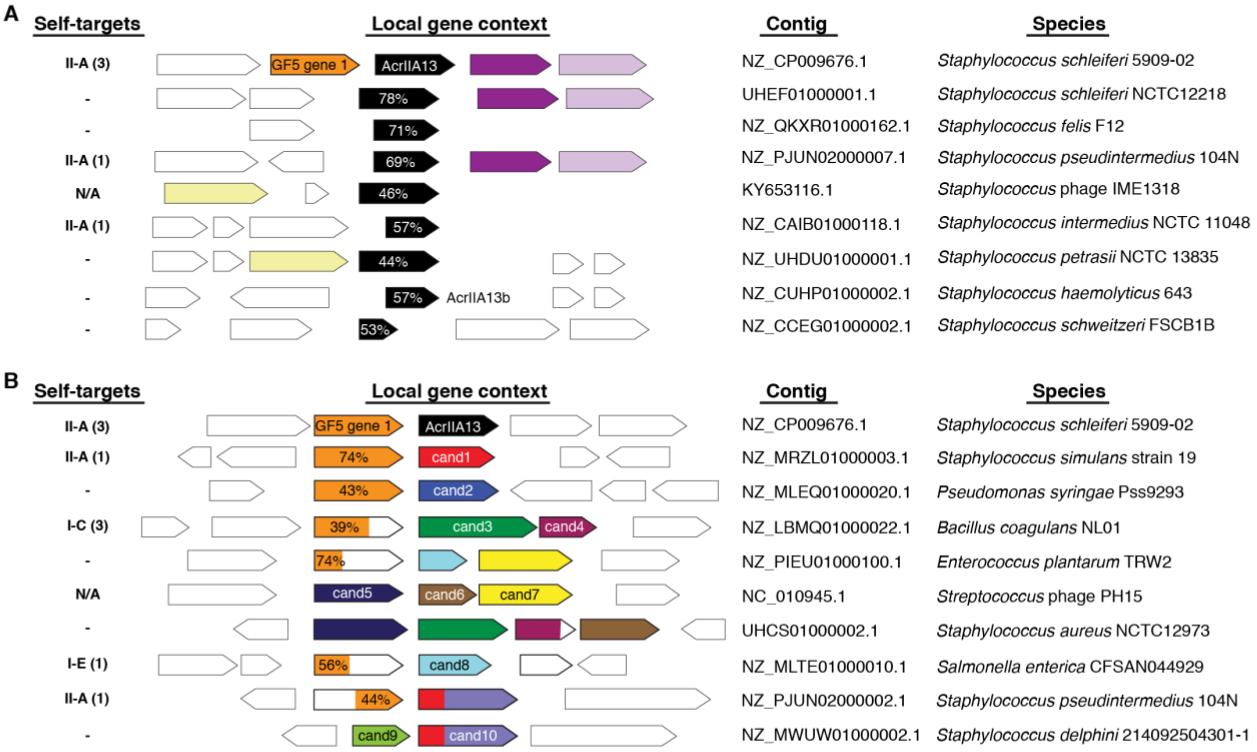


**Figure S2. Genes associated with aca8 and AcrIIA13.** (A) Using a guilt-by-association approach, we looked for genes frequently found with AcrIIA13 in the same operon. However, we only located neighboring genes that appeared to be on separate operons or genes whose homologs we had tested with TXTL during the MGE screen (Fig. 2A). Related genes share the same color. Percent identity of the AcrIIA13 homologs is shown in white text. If any self-targeting CRISPR systems were found within a species containing an AcrIIA13 homolog, they are denoted with the number of self-targets in parentheses. (B) Similar as (A), but looking for genes that co-occur with GF5 gene 1 instead. We observed 10 families of proteins that were frequently found near GF5 gene 1 or each other, example species are shown. From these 10 families, 10 proteins were chosen to be tested for Acr activity against SauCas9. Genes that share homology are colored similarly. Candidate protein 10 shares homology at its N-terminus with candidate protein 1.

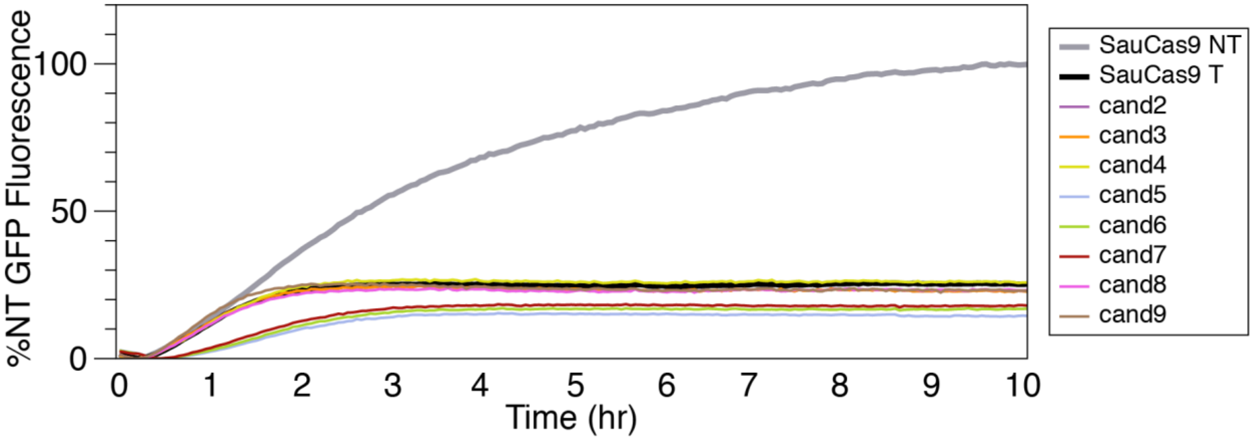


**Figure S3. Candidates 2-9 do not show Acr activity in TXTL sfGFP screen.** The TXTL screen to test individual Acrs contained the sfGFP reporter, sfGFP sgRNA plasmid, SauCas9 expression plasmid, and the indicated Acr candidate cloned into a heterologous vector. GFP fluorescence was measured every three minutes over the course of the 10 hour experiment. The non-targeting (NT) sample demonstrates high fluorescence signal after 10 hours, while the targeting (T) sgRNA shows a significant reduction in GFP expression. Acr candidates 2-9 do not show GFP expression above the targeting control, suggesting there is no Acr activity.

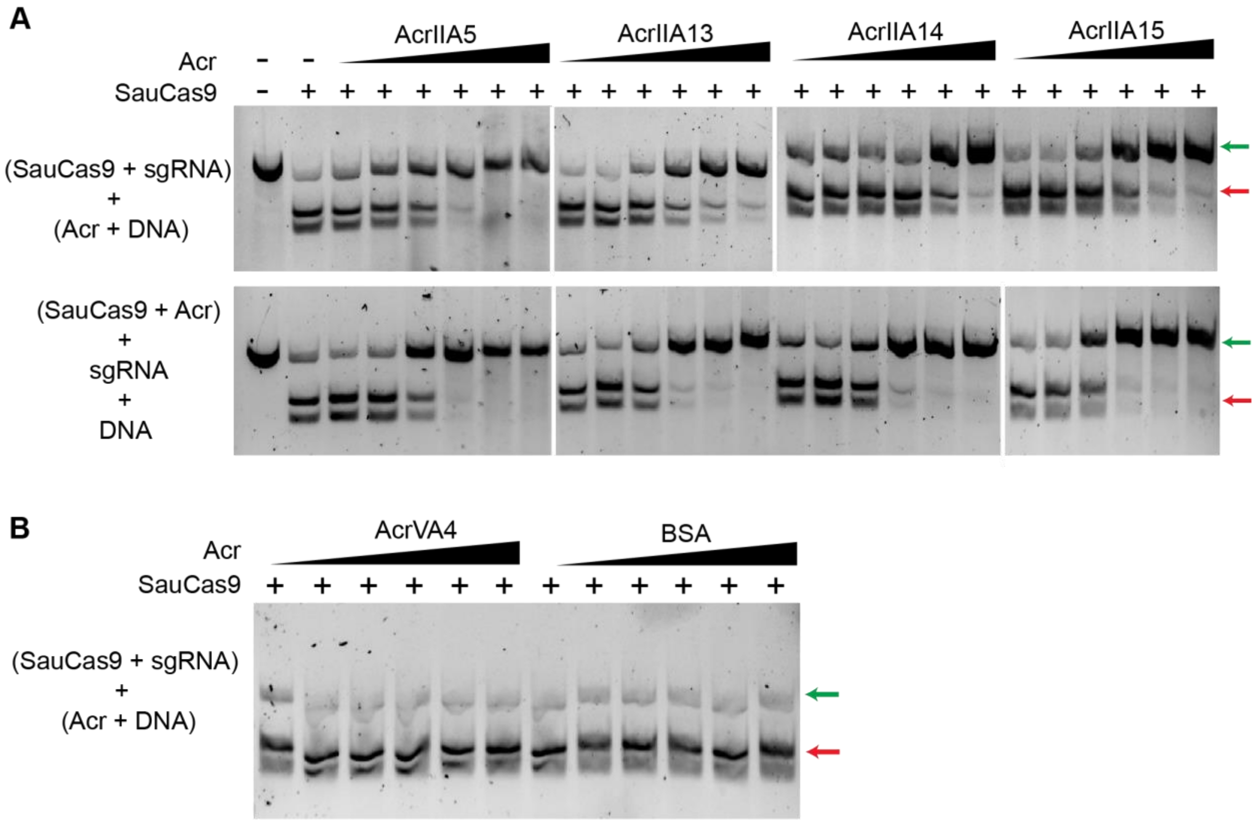


**Figure S4. AcrIIA13-15 inhibit SauCas9 cleavage activity in-vitro. (A)** Agarose gels showing cleavage of dsDNA target by SauCas9 in the presence of increasing amounts of different Acrs. Reactions were performed by adding Acrs after complexing Cas9 and sgRNA (Top panel) or prior to the addition of sgRNA (Bottom panel). The order of the addition of components are indicated besides their respective panels. These gels correspond to one of the replicates plotted in Fig. 3A. Each reaction contains 5 nM dsDNA, 100 nM SauCas9 100 nM sgRNA and the following Acr concentrations: 50 nM, 100 nM, 200 nM, 500 nM, 1000 nM and 2000 nM. **(B)** Agarose gels showing cleavage of dsDNA target by Cas9 in the presence of increasing amounts of BSA or AcrVA4 using the same range of concentrations as the Acrs in S4A. Reactions were performed by adding Acrs after complexing Cas9 and sgRNA. Uncleaved and cleaved dsDNA products are indicated by green and red arrows, respectively.

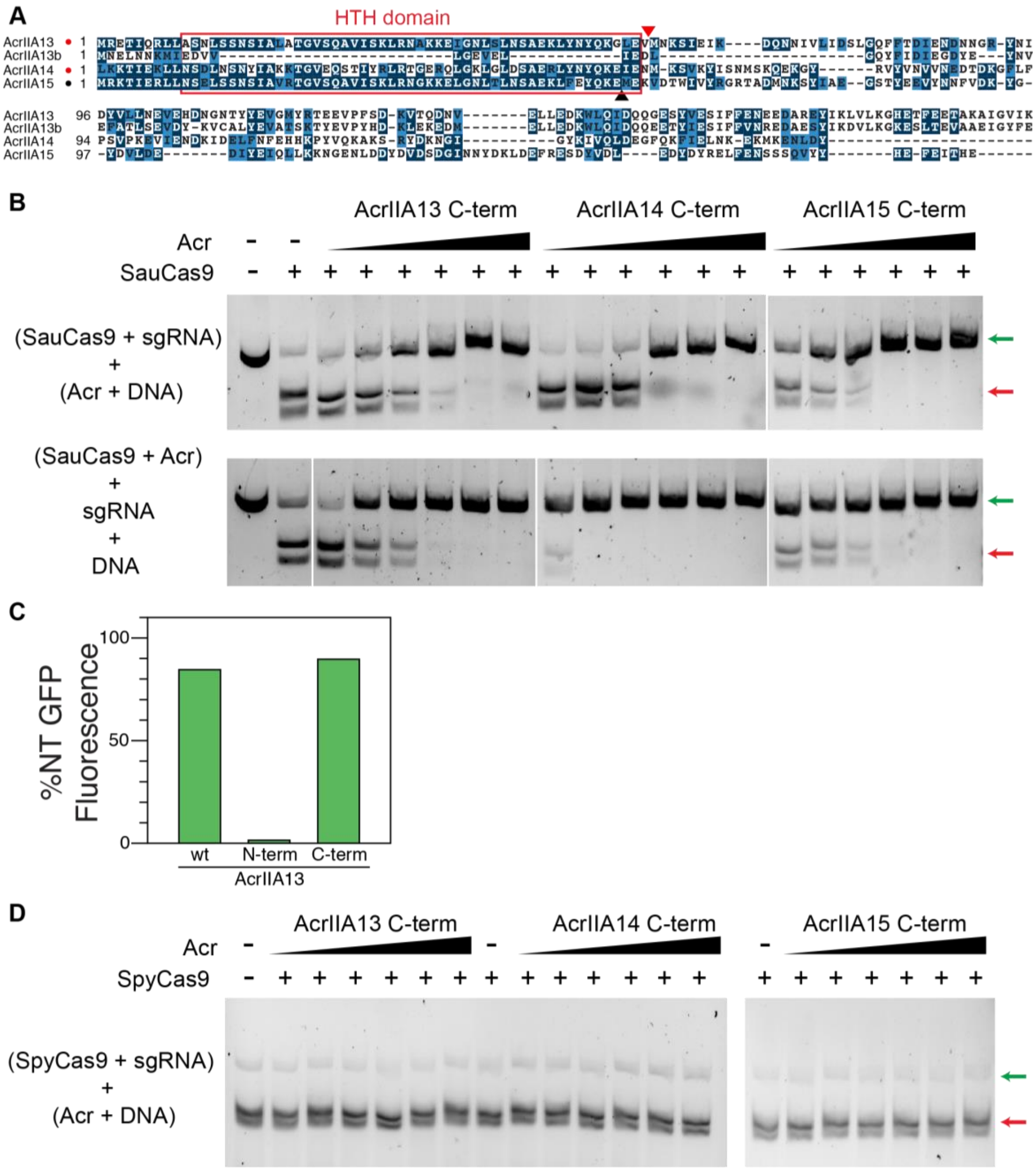


**Figure S5. N-terminal region of AcrIIA13-15 is dispensable for their Acr activity. (A)** Multiple sequence alignment for AcrIIA13-15 showing highly conserved N-terminal and divergent C-terminal. The N-terminal region of all the Acrs are predicted to contain a helix-turn-helix (HTH) motif that is highlighted by the red box. The red and black arrows mark the start codon of the C-terminal region of the Acrs. AcrIIA13b is a homolog of AcrIIA13 that is missing much of its N-terminal region. **(B)** As described in Fig. S4A except that AcrIIA13-15 C-terminal proteins were used for the assays. **(C)** TXTL results showing N-terminal region of AcrIIA13 was incapable of inhibiting SauCas9 cleavage activity while the C-terminal alone was sufficient for inhibition. Data plotted as in Fig. 2A. **(D)** Test for inhibition of SpyCas9 cleavage activity by AcrIIA13-15 C-terminal proteins. Assays were set up as described in Fig. S4A. Each reaction contains 5 nM dsDNA, 50 nM SpyCas9m 50nM sgRNA and Acr concentrations: 50 nM, 100 nM, 200 nM, 500 nM, 1000 nM and 2000 nM.

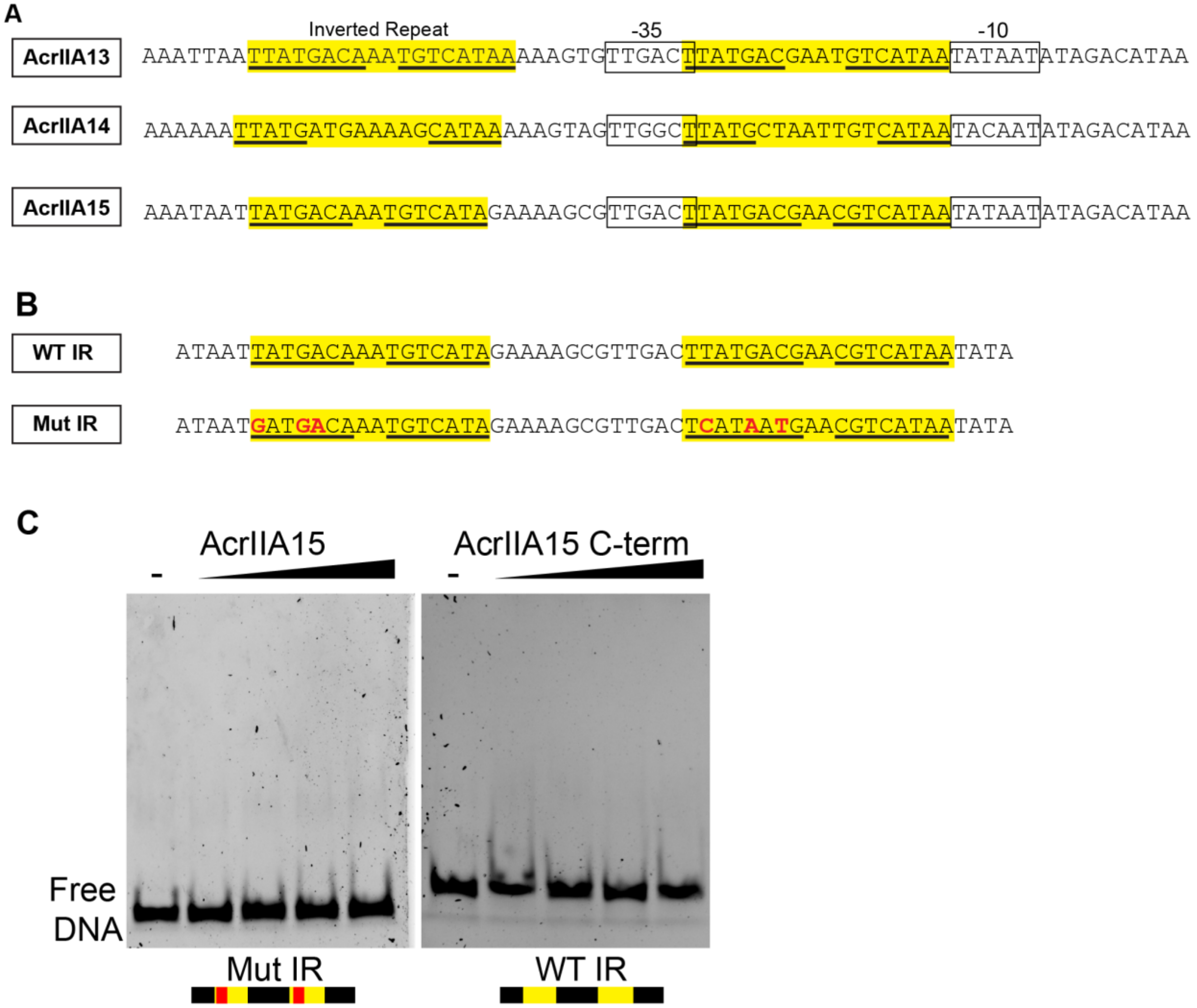


**Figure S6. AcrIIA13-15 operons contain inverted repeats (IR) proximal to their promoters. (A)** Promoter proximal sequences for AcrIIA13-15 showing the presence of two sets of inverted repeats (IR). IR segments are highlighted in yellow with the inverted repeat sequence underlined. **(B)** Sequence of the dsDNA substrates used for EMSAs in Fig. 3E and S6C. These sequences are encoded within the promoter proximal region of AcrIIA15 that were either synthesized in its wildtype form or mutated to serve as negative control. **(C)** 6% TBE polyacrylamide gels showing binding of AcrIIA15 to mutated form of the AcrIIA15 IR dsDNA substrate (Left panel) and AcrIIA15 C-term binding to WT IR DNA (Right panel).

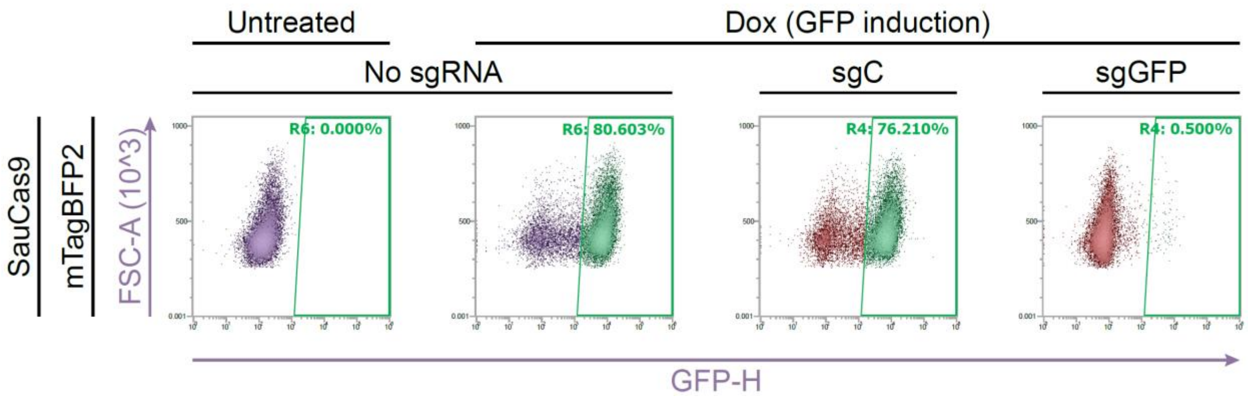


**Figure S7. Mammalian validation system to assess Acr proteins.** Assessment of genome editing inhibition efficiency by various anti-CRISPR (Acr) protein candidates or controls. HEK-RT1 genome editing reporter cells, expressing a doxycycline-inducible GFP marker, were sequentially transduced with lentiviral vectors expressing mTagBFP2 (pCF525-mTagBFP2, HygroR), SauCas9 (pCF825, PuroR), and SauCas9 sgRNAs (pCF824-sgC/sgGFP, mCherry). At day three post sgRNA transduction, the HEK-RT1 reporter cells were treated with doxycycline for 24 h, followed by flow cytometry-based quantification of GFP fluorescence. For sgRNA expressing samples, quantification was gated on the mCherry-positive population. sgC: negative control sgRNA. sgGFP: GFP-targeting sgRNA. R6: GFP-positive, live single cells. R4: GFP-positive, mCherry-positive, live single cells. GFP-H: GFP fluorescence intensity, signal height. FSC-A: forward scatter, signal area.

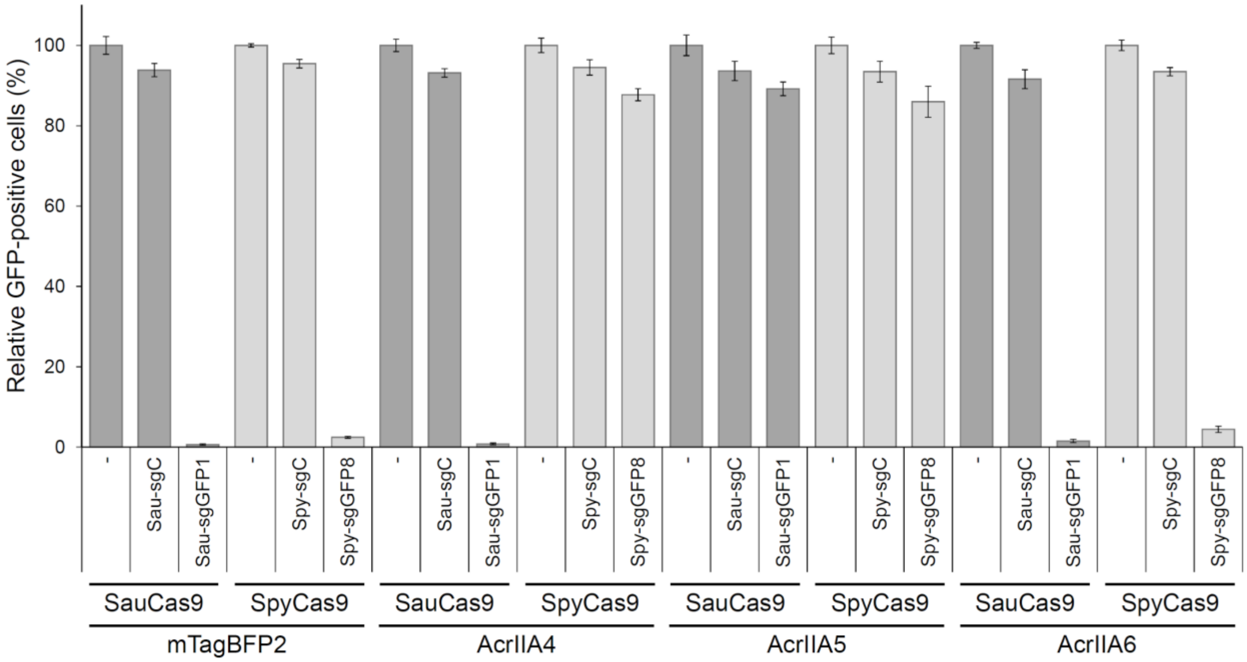


**Figure S8. Quantification of AcrIIA4, AcrIIA5, and AcrIIA6 inhibition of SauCas9.** Testing of three known Acrs for their ability to inhibit the genome editing activity of SauCas9 and SpyCas9 in human cells. HEK-RT1 genome editing reporter cells were sequentially transduced with lentiviral vectors expressing AcrIIA4, AcrIIA5, AcrIIA6, or mTagBFP2 (pCF525-AcrIIA, HygroR), SauCas9 (pCF825, PuroR) or SpyCas9 (pCF823, PuroR), and SauCas9 sgRNAs (pCF824-sgC/sgGFP, mCherry) or SpyCas9 sgRNAs (pCF820-sgC/sgGFP, mCherry). At day three post sgRNA transduction, the reporter cells were treated with doxycycline for 24 h, followed by flow cytometry-based quantification of GFP fluorescence. Error bars represent the standard deviation of triplicates. sgC: negative control guide RNAs for respective Cas9. sgGFP1/sgGFP8: GFP-targeting guide RNAs for the respective Cas9. The data shown for mTagBFP2 is partially quantification of data shown in Fig. S7.

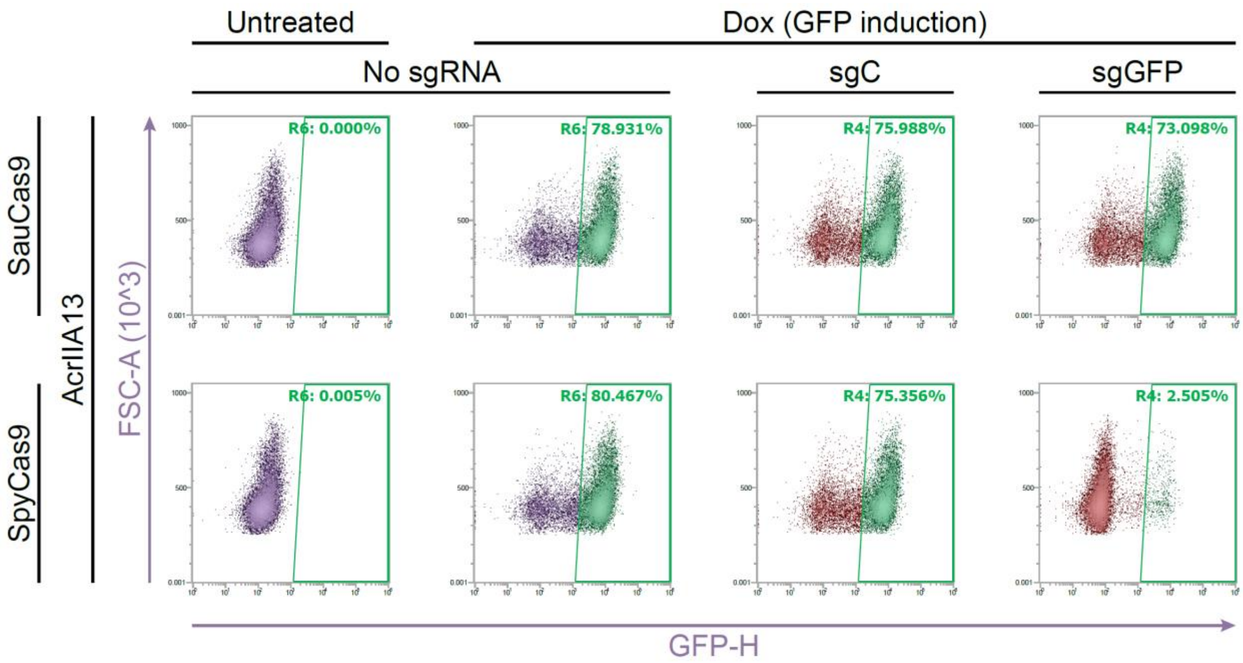


**Figure S9. AcrIIA13 potently and selectively inhibits SauCas9.** Assessment of SauCas9 and SpyCas9 genome editing inhibition efficiency by AcrIIA13 in human cells. HEK-RT1 genome editing reporter cells were sequentially transduced with lentiviral vectors expressing AcrIIA13 (pCF525-AcrIIA, HygroR), SauCas9 (pCF825, PuroR) or SpyCas9 (pCF823, PuroR), and SauCas9 sgRNAs (pCF824-sgC/sgGFP, mCherry) or SpyCas9 sgRNAs (pCF820-sgC/sgGFP, mCherry). At day three post sgRNA transduction, the reporter cells were treated with doxycycline for 24 h, followed by flow cytometry-based quantification of GFP fluorescence. sgC: negative control sgRNAs for respective Cas9. sgGFP: GFP-targeting sgRNAs for the respective Cas9. R6: GFP-positive, live single cells. R4: GFP-positive, mCherry-positive, live single cells. GFP-H: GFP fluorescence intensity, signal height. FSC-A: forward scatter, signal area.

**Table S1. List of expected lethal self-targeting *Staphylococcus* genomes obtained with Self-Target Spacer Searcher (STSS).** 11,910 *Staphylococcus* assemblies were downloaded from NCBI and run using STSS {Watters:2018kv}, resulting in 99 total instances of self-targeting. Self-targets marked as ambigious type II CRISPR systems were further manually curated as type II-A by observation of similar identity to known type II-A Cas9 proteins as well as homology of a gene between Cas2 and the array to known Csn2 genes, which are poorly captured in the Hidden Markov Models used in STSS. Type II-A CRISPR systems with PAM sequences within one indel or mutation of the known 3′-NNGRR(T) were filtered from the list of all self-targeting instances and also listed separately.

See Table_S1.xlsx

**Table S2. Filtered List of *Staphylococcus* II-A self-targeting instances within one indel/mutation of the NNGRR(T) PAM.** The list of genomes containing type II-A self-targeting instances was filtered by comparing the bases downstream of the aligned target sequence to the known SauCas9 PAM sequence, NNGRR(T). Self-targeting instances were kept if the sequences downstream of the target matched within one indel or mutation of SauCas9 PAM sequence (final T optional), given these sequences would be most likely to yield a lethal self-target and thus be more likely to contain an Acr gene. The PAM filtered list resulted in 12 genomes containing type II-A CRISPR self-targeting. To narrow the search further, the list was sorted based on the similarity of each system's Cas9 to SauCas9 (CCK74173.1 from S. aureus subsp. aureus M06/0171). We also used PHASTER {Arndt:2016hm}and Islander {Hudson:2015hs} to determine if any of the self-targets occurr within prophages. Self-targets found in prophages with no target sequence mutations and a bona fide PAM sequence are hallmarks of the scenario described in Fig. 1A, where protospacers are acquired from a Acr-carrying phage before the phage integrates to create the self-targeting spacer.

| Identity to SauCas9 | Cas9 Accession | Species/Strain | Spacer Sequence | Target Sequence | PAM Region (Downstream) | Repeat Mutations (vs. Consensus) | Target Location/Predicted Gene(s) | In MGE? |
|---|---|---|---|---|---|---|---|---|
| 100% | WP_053019794.1 | Staphylococcus haemolyticus strain W_75 | TAATTCCAACTTAACAACCAATCCAAATCAT | Perfect match | TAAAGCGAT | None | Between WP_053020048.1, Replication and maintenance protein & locus tag: AK682_RS12120, CDS | No MGEs predicted |
| 100% | WP_053019794.1 | Staphylococcus haemolyticus strain W_139 | TAATTCCAACTTAACAACCAATCCAAATCAT | Perfect match | TAAAGCGAT | None | Between WP_053020048.1, Replication and maintenance protein & locus tag: AK682_RS12120, CDS | No MGEs predicted |
| 86% | WP_105980293.1 | Staphylococcus simulans strain 19 | TACGAATCCCAGCACGTATAATTCAAGTTT | .g........................... | TGAAATTTT | None | WP_105980054.1, hypothetical protein | N |
| 80% | WP_105994700.1 | Staphylococcus simulans strain 116 | ATGTTTTACAAGAAGAATTAGCACCTGATC | Perfect match | TAAAAACAA | None | WP_009384743.1, hypothetical protein | No MGEs predicted |
| 62% | WP_050331073.1 | Staphylococcus schleiferi strain 2317-03 | ACGGTGGAAAACTTGAATTTGAAGCTGATCA | Perfect match | AGAAGTGAA | None | WP_050329766.1, PTS glucose transporter subunit IICBA | N |
| | | Staphylococcus schleiferi strain 2317-03 | AATGGCTTACATTAACAAATTCAACGAAAT | Perfect match | GGAAAATGC | None | WP_050331546.1, Rha family transcriptional regulator | Y |
| | | Staphylococcus schleiferi strain 5909-02 | AATGGCTTACATTAACAAATTCAACGAAAT | Perfect match | GGAAAATGC | None | WP_050337612.1, Rha family transcriptional regulator | Y |
| 58% | WP_044361501.1 | Staphylococcus microti strain NCTC13832 | TCATGTTTTTAGGGTCGTTTGCTGCACCAA | ..........................g. | TAGGGGCAT | None | WP_044359341.1, CHAP domain-containing protein | N |
| 27% (69% coverage) | WP_096665615.1 | Staphylococcus delphini strain 14S03314-1 NODE_12_length_102660_cov_47.2899_ID_23 | TTGAGTGTCAACGTTGTAAGTCGAAAGGA | Perfect match | AGGGTAGCT | Upstream repeat mutated: ……………….a….. | WP_096649624.1, alpha/beta hydrolase | Y |
| | | Staphylococcus delphini strain 14S03314-1 NODE_12_length_102660_cov_47.2899_ID_23 | TGTTTCAGACTCACCAATCATTGCGACATA | Perfect match | TGGCGCTCT | Downstream repeat mutated: ……………….a….. | WP_096665888.1, phage tail protein | Y |
| 27% (69% coverage) | WP_096598476.1 | Staphylococcus delphini strain 15S02591-1 NODE_21_length_38160_cov_72.6355_ID_41 | ACGTTTATTATCTGAACGCCGTTTTTGCT | Perfect match | CGGGTTTTT | None | Between WP_096597505.1, hypothetical protein & WP_096597503.1, hypothetical protein | Y |
| 26% (67% coverage) | WP_096601671.1 | Staphylococcus intermedius strain 14S03297-1 NODE_2_length_897250_cov_66.0175_ID_3 | TCCTAGACCTAATCATAGGTTTGTGGAGG | Perfect match | GGGGGAAGA | None | WP_019168966.1, RusA family crossover junction endodeoxyribonuclease | No MGEs predicted |

| | | | | | | | | |
|---|---|---|---|---|---|---|---|---|
| | | Staphylococcus intermedius strain 14S03299-1 NODE_2_length_897243_cov_78.3361_ID_3 | ATATTTTTAGCGCCTATGTTCCCTTTAGGT | Perfect match | GCTAGTATC | None | WP_096601956.1, MFS transporter | No MGEs predicted |
| 27% (53% coverage) | WP_096536567.1 | Staphylococcus intermedius strain 14S03299-1 NODE_2_length_897243_cov_78.3361_ID_3 | TCCTAGACCTAATCATAGGTTTGTGGAGG | Perfect match | GGGGGAAGA | None | WP_019168966.1, RusA family crossover junction endodeoxyribonuclease | No MGEs predicted |

**Table S3. Genomic amplicons tested for Acr activity.** The predicted MGEs found in *S. schleiferi* strain 5909-02 and *S. haemolyticus* strain W75 were broken up into 27 ~10 kb genomic fragment (GF) amplicons in total to screen for Acr activity. We were unable to amplify one of the *S. haemolyticus* fragments, thus it was not screened. The three self-targets in *S. schleiferi* strain 5909-02 can be found distributed across three genomic fragment amplicons. The self-target for *S. haemolyticus* is not captured by the amplicons because it falls in a contig that is too short for reliable MGE prediction, and does not contain any complete genes. The Cas9 amplicons used for GF screening (Fig. 2A) are also indicated.

| Contig | Self-target? | Organism | Forward primer | Reverse primer | Len (kb) | Name |
|---|---|---|---|---|---|---|
| NZ_CP009676.1 | | *S. schleiferi* strain 5909-02 | ggcgcgacctacaaagtgc | ggtctaatttttccaccgttttactcctc | 9.9 | GF1 |
| | | | gaggaggagtaaaacggtggaaaaattag | tgtggttgtattttttatggtgttcaact | 9.7 | GF2 |
| | | | gaattgaaaacttaaaaactcggtgaagttgaacac | gtgttctgccgtttactattactgtatcg | 9.7 | GF3 |
| | | | aatcaagattcaaaccgggaggtgc | tacagtatacataaacttcactcctctccaatg | 9.8 | GF4 |
| | | | gaaaaaattaaagaatacattggagaggagtgaag | aaagaatattaaaataaattttaagaagaatgtttaaagaaatgcgg | 7.7 | GF5 |
| | | | caagataagaaaaaacttgtttcaatcatttaaaaaagg | ggtgatgtaatgcaaaaaaatgaactcc | 11.4 | GF6 |
| | | | catttttttgcattacatcaccattttctttaaacatatc | agttaaaacaaagggggattcaaatgatg | 9.0 | GF7 |
| | | | atgagaataatcatactcataattacaacctccc | gatgaggtgtttaaatgatttttaaaagcaaag | 10.4 | GF8 |
| | Y | | ctttgcttttaaaaatcatttaaacacctcatc | gaaaggggttaatataattgattgctcacga | 9.3 | GF9 |
| | | | gagcaatcaattatattaacccctttccttaaaac | tgaatctaaacggtgcggtgacattta | 11.5 | GF10 |
| | Y | | gtcgtcattaaatgtcaccgcacc | ttcagaaaaattataccagaaaggagataaaaaaatggc | 7.9 | GF11 |
| | Y | | atgtagccatttttttatctcctttctgg | acgaaggagtttttctcatatgaattttaataacc | 10.8 | GF12 |
| | | | gcgagactacaaaagttagaatcactg | cacactatgtcactttacttgctatatgag | 5.5 | GF13 |
| | | | tgtgaattgctgttatgtttttaagaagcttatc | catctcagcacttcgtctttatttttgc | 12.3 | GF14 |
| | | | atttgtcatttattacgaatcccggtggg | gagtacataagagatcgaacagaagctag | 12.2 | GF15 |
| | | | gcaagtttatgttagacatttttaaattttcg | aggaagaagggctgacatagtctaac | 6.0 | Cas genes |
| NZ_CUFQ01000045.1 | | *S. haemolyticus* strain W_75 | cagcctttcgtttaatttataagttaatgatagc | agacatcaccgtggtgcaac | 10.5 | GF16 |
| NZ_CUFQ01000031.1 | | | cttatccaagaactttatgtcccggactc | catcaatctcaccgtctcttaaggtatc | 13.1 | GF17 |
| NZ_CUFQ01000018.1 | | | ggatagacggactacccgtaatattacac | gtctctagtgctaaatctttttttcatttcttaattcct | 8.0 | GF18 |
| NZ_CUFQ01000018.1 | | | cagcagtctctttgattgtcaaaaacat | cgtttgataatacatccatacttctctcacac | 10.0 | GF19 |
| NZ_CUFQ01000008.1 | | | ccactacaatcgcaattgctttaaaatc | ttgaagatggtgacgaacgcatacaaag | 12.4 | GF20 |
| NZ_CUFQ01000008.1 | | | ttgaagatggtgacgaacgcatacaaag | gttttaggtgattttgctaatggtagacg | 12.5 | GF21 |
| NZ_CUFQ01000005.1 | | | gttttaggtgattttgctaatggtagacg | tccgtaagagtacctttaatcgatagtcattg | 8.8 | GF22 |
| NZ_CUFQ01000003.1 | | | aagagttacgcaattattagagatggaatcc | aatgacacctcttaatcattatttatattaaattattatagcaaatgc | 7.2 | GF23 |
| NZ_CUFQ01000005.1 | | | catgacctcttaaattcatacttaaagaccc | gtgatagtgatgtagttcatatacatcaatggg | 8.7 | GF24 |
| NZ_CUFQ01000005.1 | | | gcatcaattatttcagcaagtattattggc | cgagcaagtatacagagcatttaaaatcatg | 9.7 | GF25 |
| NZ_CUFQ01000004.1 | | | cggagacgttaaatcaaaagcagattttg | cggtcttactttgggttattattgctc | 10.3 | - |
| NZ_CUFQ01000006.1 | | | atccttctggcgatgtcttaacc | accacattcctttaacctaattgttgtttc | 6.8 | GF26 |
| NZ_CUFQ01000030.1 | | | gagtactaaaacctctagcatcattataaatcctc | tcaccattgcctctttagttatttcatgtg | 5.8 | Cas genes |

**Table S4. Acr candidates identified using guilt-by-association approach.** A summary of the 10 different candidates tested for Acr activity in TXTL identified from the guilt-by-association approach using GF5 gene 1 (Fig. S2B, S3, Fig. 2E). Candidate proteins 1 and 10 were found to inhibit SauCas9. Proteins marked with an asterisk were not predicted correctly on the NCBI database and therefore do not have an accession number, or the accession number contains a shortened form of the protein. The proteins found in GF5 used to seed the guilt-by-association search are also listed. Candidate 1 was tested in TXTL using a methionine start codon, replacing the wildtype leucine.

| Name | Accession | Amino Acid Sequence |
|---|---|---|
| GF5 gene 1 | WP_050337584.1 | MNKSNLFKTAWEIAKNGASKFGGSCKDYFAESLKMAYKGIVLAEAEIEIPAWIVRK NVGNVYVVEKSVLSVKRETEKALLIHADGKFGAFDFWTPKSVLKTVNVVDYTQATI TVENEMVKAMDNHFELVKKAKALGIKGIRSNMKSSTLRKKIAEVA |
| GF5 gene 2 (AcrIIA13) | WP_050337628.1 | MEVMNKSIEIKDQNNIVLIDSLGQFFTDIENDNNGRYNIDYVLLNEVEHDNGNTYYE VGMYRTEEVPFSDKVTQDNVELLEDKWLQIDQQGESYVESIFFENEEDAREYIKLVL KGHETFEETAKAIGVIK |
| AcrIIA13b | WP_053038109.1 | MNELNNKMIEDVVLGEVELIEDLGQYFIDIEGDYEYNVEFATLSEVDYKVCALYEV ATSKTYEVPYHDKLEKEDMKLFYDKWLEKDQQEETYIESVFFVNREDAESYIKDVL KGKESLTEVAAEIGYFE |
| candidate 1* (AcrIIA14) | - | LKKTIEKLLNSDLNSNYIAKKTGVEQSTIYRLRTGERQLGKLGLDSAERLYNYQKEIE NMKSVKYISNMSKQEKGYRVYVNVVNEDTDKGFLFPSVPKEVIENDKIDELFNFEH HKPYVQKAKSRYDKNGIGYKIVQLDEGFQKFIELNKEKMKENLDY |
| candidate 2 | WP_103363596.1 | MNDLENPLEIVYTSREAAELYGLSENTVTQWCNRGRFTDEEARKSSKVWLVTKKG MERLTNKEEKKLKIALSTNPTVEKYLGLALEHNSPFEYNHTTFENIKRLYDFVVDNS EEGFIFEEVLKEVIDRAAVQGGPSVSYELGSHETKSGHAESISFDYDSEYEEDEDTYT STIIF |
| candidate 3 | WP_052749681.1 | MKRNDVDLTGRTFYDLNVKCRHTEKLYNETAWVCKCLACGNITYATTSQLLHGKK KSCGCRRKKTPPNALDLTNKQFGKLKVVERAGISKDGRALWLCHCKCGNTIVTNAT SLRIGDVKSCGKCLVDERIKHAREELMTNKTIDGVPVPLLTKKVRSDSGTGHKGVQ RRVRKGKEKFEAYITVKGKRKYIGTYSKLDDAIKARKDAEKEYFEPYIKALKEREKE |
| candidate 4 | WP_046721438.1 | MEKQRVNMDINKELWKKVGIRAIELNLQKKEIVELALEKFLQEGNKMAKVFYEDIL VGEVLTNRSLTVEEALEAIGFDEAKFIEEQGFDDIDPNDFRVEY |
| candidate 5 | YP_001974381.1 | MINVNNTNSAELENVVVDGQLCLSASKRQPVWVAEIVGTHPVYKLDRKFIDEDESG AGWKAWNLEENKIYCINPSANKKDQYFVVLVDGALNELTKGQVEEMVK |
| candidate 6 | YP_001974382.1 | MLVGVSQKNDFGWSAQVVKFPNLESAEAWLNKEQFDFRDRYIFDDEEEALNHLKEI KSVSWAKEALKDADTLTLLDNGEFDIEMSDSYIYKMMTR |
| candidate 7 | YP_001974383.1 | MKFELFKELYNEALGIASLDLFVAERGWQDWMEDYDADDVVKILTNIYYLANNPL KDTRKVSRAEFSRQYNIPNRTLQDWDLGNRNAPEYVKLLLDFAQFTNR |
| candidate 8 | WP_070801998.1 | MNNTEKLMAVGKLVYGDNWQSPISRDIGVDSRTIRYALKGEREINHLSSRLKEALE QKIEKLKSAIEIINSDKMSGDDVDVDIISNIVDGYEYSDEQYKKAAFDEINNVVCADT WLSDLDSIARKWSKY |
| candidate 9 | PCF60931.1 | MFVTKKELKDLNVKEVFESGKNFISIKDGRHIIYRVNGKYLVSRDDKLYPETRIPKYI QRKLV |
| candidate 10 (AcrIIA15) | WP_142302263.1 | MRKTIERLLNSELSSNSIAVRTGVSQAVISKLRNGKKELGNLTLNSAEKLFEYQKEM EKVDTWIVYRGRTADMNKSYIAEGSTYEEVYNNFVDKYGYDVLDEDIYEIQLLKKN GENLDDYDVDSDGINNYDKLDEFRESDYVDLEDYDYRELFENSSSQVYYHEFEITH E |

**Table S5. DNA sequences used in TXTL.** Exact oligonucleotide sequences are shown, while plasmid sequences are described. All Acr expression plasmids were cloned into the same vector driven by pTet. The plasmids in the table marked with an asterisk can be found on Addgene with #-##.

| Name | Sequence | Description |
|---|---|---|
| Chi6.fwd | TCACTTCACTGCTGGTGGCCACTGCTGGTGGCCACTGCT<br>GGTGGCCACTGCTGGTGGCCACTGCTGGTGGCCACTGCT<br>GGTGGCCA | Annealed with Chi6.rev to prevent dsDNA degradation in TXTL |
| Chi6.rev | TGGCCACCAGCAGTGGCCACCAGCAGTGGCCACCAGCA<br>GTGGCCACCAGCAGTGGCCACCAGCAGTGGCCACCAGC<br>AGTGAAGTGA | Annealed with Chi6.fwd to prevent dsDNA degradation in TXTL |
| pKEW002 | p15A - CmR - TetR - SauCas9 - terminator | SauCas9 expression regulated by pTet |
| sfGFP plasmid | ColE1 - AmpR - J23119 - sfGFP - terminator | sfGFP reporter plasmid |
| pKEW037 | ColE1 - AmpR - LacI - $P_{Lac}$ - GFP sgRNA - terminator - araC<br>sgRNA sequence (spacer in lowercase):<br>actggagttgtcccaattcttgGTTTTAGTACTCTGGAAACAGAATCT<br>ACTAAAACAAGGCAAAATGCCGTGTTTATCTCGTCAACT<br>TGTTGGCGAGATTTTT | sgRNA plasmid targeting sfGFP |
| pKEW278* | ColE1 - AmpR - TetR - $P_{tet}$ - Acr - candidate | AcrIIA13 under pTet |
| pKEW312* | | AcrIIA13a under pTet |
| pKEW277* | | GF5 cand1 under pTet |
| pKEW345* | | Candidate 1 under pTet (AcrIIA14) |
| pKEW346* | | Candidate 2 under pTet |
| pKEW347* | | Candidate 3 under pTet |
| pKEW348* | | Candidate 4 under pTet |
| pKEW349* | | Candidate 5 under pTet |
| pKEW350* | | Candidate 6 under pTet |
| pKEW351* | | Candidate 7 under pTet |
| pKEW352* | | Candidate 8 under pTet |
| pKEW372* | | Candidate 9 under pTet |
| pKEW373* | | Candidate 10 under pTet (AcrIIA15) |
| pKEW332* | | AcrIIA13 N-terminus under pTet |
| pKEW331* | | AcrIIA13 C-terminus under pTet |

**Table S6. DNA/RNA sequences used for biochemistry.**

| Name | Sequence | Description |
|---|---|---|
| SauCas9 EGFP sgRNA | gcaagggcgaggagctgttcacGTTTTAGTACTCTGGAAACAGAATC<br>TACTAAAACAAGGCAAAATGCCGTGTTTATCTCGTCAAC<br>TTGTTGGCGAGATTTTT | SauCas9 sgRNA that targets EGFP |
| WT IR DNA P1 | ATAATTATGACAAATGTCATAGAAAAGCGTTGACTTATG<br>ACGAACGTCATAATATA | Used for DNA EMSA in Fig 3E |
| WT IR DNA P2 | TATATTATGACGTTCGTCATAAGTCAACGCTTTTCTATGA<br>CATTTGTCATAATTAT | Used for DNA EMSA in Fig 3E |
| Mut IR DNA P1 | ATAATGATAGCAAATGTCATAGAAAAGCGTTGACTCAT<br>AATGAACGTCATAATATA | Used for DNA EMSA in Fig S6C |
| Mut IR DNA P2 | TATATTATGACGTTCATTATGAGTCAACGCTTTTCTATGA<br>CATTTGCTATCATTAT | Used for DNA EMSA in Fig S6C |
| FAM_EGFP_P1 | tggtgagcaagggcgaggagctgttcaccggggt | Target dsDNA (T) used for DNA EMSA in Fig 3C |
| FAM_EGFP_P2 | accccggtgaacagctcctcgcccttgctcacca | Target dsDNA (T) used for DNA EMSA in Fig 3C |
| FAM_ctrl_P1 | tacgtccaggagcgcaccatcttcttcaaggacg | Control non-target dsDNA (NT) used for DNA EMSA in Fig 3C |
| FAM_ctrl_P2 | cgtccttgaagaagatggtgcgctcctggacgta | Control non-target dsDNA (NT) used for DNA EMSA in Fig 3C |
| Cleavage target DNA | cccttgaacctcctcgttcgaccccgcctcgatcctccctttatccagccctcactccttctctag<br>gcgccggaattgaagatctatcgattctagattcgagtttactccctatcagtgatagagaacgt<br>atgaagagtttactccctatcagtgatagagaacgtatgcagactttactccctatcagtgatag<br>agaacgtataaggagtttactccctatcagtgatagagaacgtatgaccagtttactccctatca<br>gtgatagagaacgtatctacagtttactccctatcagtgatagagaacgtatatccagtttactcc<br>ctatcagtgatagagaacgtataagctttaggcgtgtacggtgggcgcctataaaagcagagc<br>tcgtttagtgaaccgtcagatcgcctggagcaattccacaacacttttgtcttataccaactttcc<br>gtaccacttcctaccctcgtaaagtcgagcttgcgttggatccaccatggtgagcaagggcga<br>ggagctgttcaccggggtggtgcccatcctggtcgagctggacggcgacgtaaacggccac<br>aagttcagcgtgtccggcgagggcgagggcgatgccacctacggcaagctgaccctgaag<br>ttcatctgcaccaccggcaagctgcccgtgccctggcccaccctcgtgaccaccctgaccta<br>cggcgtgcagtgcttcagccgctaccccgaccacatgaagcagcacgacttcttcaagtccg<br>ccatgcccgaaggctacgtccaggagcgcaccatcttcttcaaggacgacggcaactacaa<br>gacccgcgccgaggtgaagttcgagggcgacaccctggtgaaccgcatcgagctgaagg<br>gcatcgacttcaaggaggacggcaacatcctggggcacaagctggagtacaactacaacag<br>ccacaacgtctatatcatggccgacaagcagaagaacggcatcaaggtgaacttcaagatcc<br>gccacaacatcgaggacggcagcgtgcagctcgccgaccactaccagcagaacacccccc<br>atcggcgacggccccgtgctgctgcccgacaaccactacctgagcacccagtccgccctga<br>gcaaagaccccaacgagaagcgcgatcacatggtcctgctggagttcgtgaccgccgccg<br>GGATCACTCTCGGCATGGACGA | dsDNA target used for cleavage assays |

**Table S7. Lentiviral vector sequences.** The complete sequences of all the lentiviral vectors used for stable transduction and expression of SauCas9 or SpyCas9, respective sgRNAs, and Acrs or mTagBFP2 are listed with a brief description of their key features. Plasmids XXX, XXX are also available on Addgene as ###, ###, respectively.

See Table_S7.xlsx
